# CUB and Sushi Multiple Domains 1 (CSMD1) opposes the complement cascade in neural tissues

**DOI:** 10.1101/2020.09.11.291427

**Authors:** Matthew L. Baum, Daniel K. Wilton, Allie Muthukumar, Rachel G. Fox, Alanna Carey, William Crotty, Nicole Scott-Hewitt, Elizabeth Bien, David A. Sabatini, Toby Lanser, Arnaud Frouin, Frederick Gergits, Bjarte Håvik, Chrysostomi Gialeli, Eugene Nacu, Anna M. Blom, Kevin Eggan, Matthew B. Johnson, Steven A. McCarroll, Beth Stevens

## Abstract

Schizophrenia risk is associated with increased gene copy number and brain expression of *complement component 4* (*C4*). Because the complement system facilitates synaptic pruning, the *C4* association has renewed interest in a hypothesis that excessive pruning contributes to schizophrenia pathogenesis. However, little is known about complement regulation in neural tissues or whether such regulation could be relevant to psychiatric illness. Intriguingly, common variation within *CSMD1*, which encodes a putative complement inhibitor, has consistently associated with schizophrenia at genome-wide significance. We found that Csmd1 is predominantly expressed in the brain by neurons, and is enriched at synapses; that human stem cell-derived neurons lacking CSMD1 are more vulnerable to complement deposition; and that mice lacking Csmd1 have increased brain complement activity, fewer synapses, aberrant complement-dependent development of a neural circuit, and synaptic elements that are preferentially engulfed by cultured microglia. These data suggest that CSMD1 opposes the complement cascade in neural tissues.

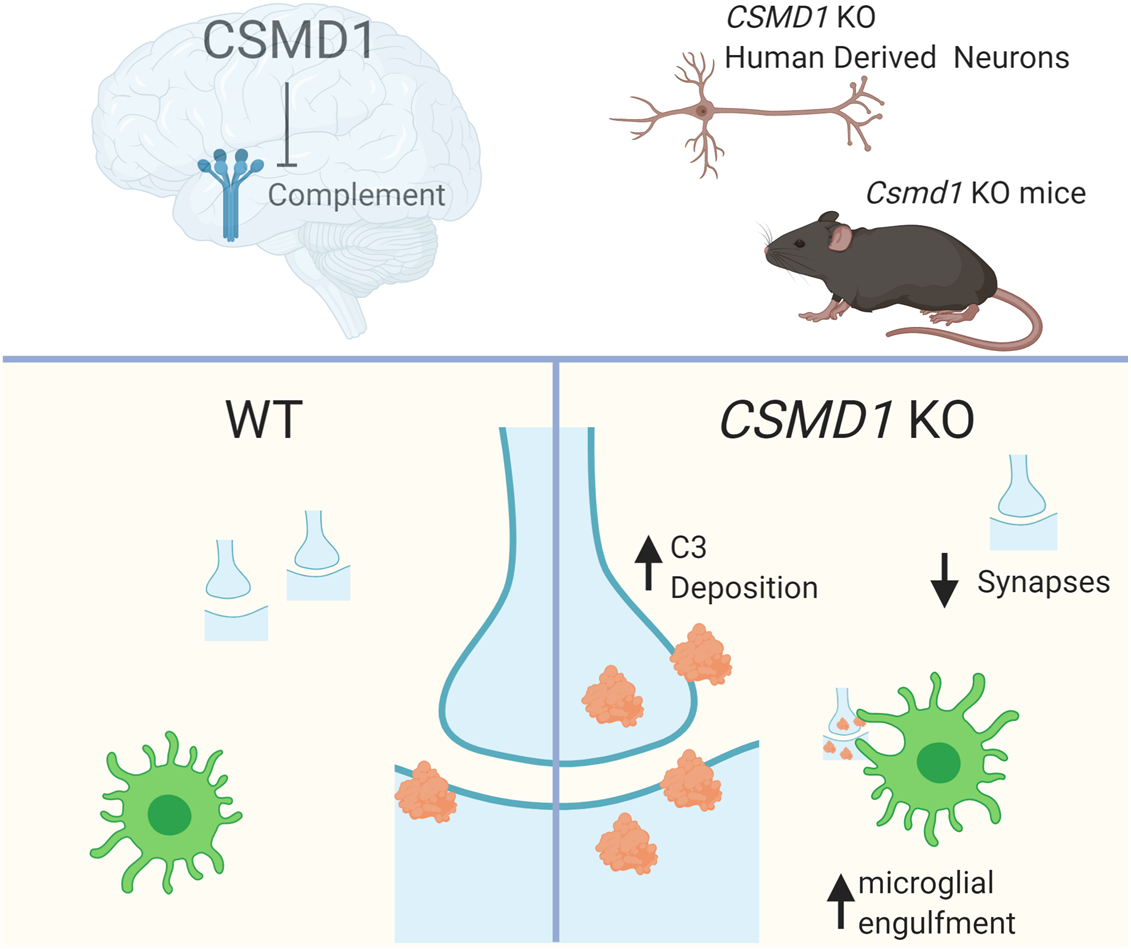
Graphic Abstract. Our findings support a model in which CSMD1 opposes actions of the complement cascade in neural tissues (top left). We investigated two models in which Csmd1 was genetically ablated: human cortical neurons derived from embryonic stem cells, and a back-crossed C57bl6-Tac mouse line (top right). Csmd1 is normally expressed by neurons and present at synapses where it can protect them from complement (bottom left); in the absence of Csmd1 (bottom right), we find more deposition of complement (on cultured human cortical neurons and in the mouse visual system), reduced numbers of synapses (in the mouse visual system), and synaptic fractions that are more readily engulfed by microglia (*ex vivo*). Created with BioRender.com.

## Introduction

Schizophrenia is a heritable, polygenic psychiatric disorder characterized by delusions, hallucinations, flattened affect, and progressive, treatment refractive cognitive impairment (Elvevåg and Goldberg, 2000; McCarroll and Hyman, 2013). Many hypotheses about disease pathogenesis have been proposed. Large-scale human genetics studies have identified associations of schizophrenia risk to common variants. One of the genes associated with schizophrenia at genome wide significance is *CSMD1* (CUB and Sushi Multiple Domains 1), which encodes a ∼388kDa type-I transmembrane protein sharing structural synteny with known complement inhibitors (Schizophrenia Psychiatric Genome-Wide Association Study, 2011; Schizophrenia Working Group of the Psychiatric Genomics, 2014; Lam et al., Nat Genet 2019). CSMD1 is named for its large extracellular domain which consists of multiple CUB and Sushi domains, protein motifs that are conserved in regulators of the complement cascade (Escudero-Esparza et al., 2013; Havik et al., 2011; Kraus et al., 2006). Consistent with this structural synteny, recombinant short fragments of the tandem-sushi region of CSMD1 inhibit activation of the complement cascade *in vitro*, and reducing endogenous CSMD1 levels in a cancer cell line results in increased cell-surface complement deposition (Escudero-Esparza et al., 2013; Kraus et al., 2006). However, the extent to which CSMD1 can regulate complement activity in the brain, as well as which brain cells and surfaces endogenously host CSMD1, are unknown.

The complement cascade involves a set of secreted, fluid-phase innate immune molecules; these molecules are repurposed in the developing or diseased nervous system to label synapses for elimination (pruning) by microglia, the brain’s resident immune cells (Hong et al., 2016; Lehrman et al., 2018; Schafer et al., 2012; Stephan et al., 2012; Stevens et al., 2007; Dejanovic et al., 2018; Wu et al., 2019). Genetic evidence for the relevance of the complement cascade to schizophrenia comes from the finding that structural variation in the *complement component 4* (*C4*) genes within the MHC locus underlies the region’s strong genome-wide association signals with schizophrenia risk (Sekar et al., 2016; Kamitaki et al., 2020). The human genome contains two paralogous C4 genes, *C4A* and *C4B*. Schizophrenia risk increases with gene copy number of *C4A*, and increased *C4A* gene copy number results in increased C4A mRNA expression in human brain tissue. Mouse studies have further shown that changes in C4 expression levels alter C3 deposition and synapse number in frontal cortex and visual thalamus (Comer et al., 2020; Sekar et al., 2016). Combined, these observations suggest that enhanced activity of the complement cascade contributes to schizophrenia pathogenesis.

One mechanism by which over-activation of the complement cascade could contribute to schizophrenia pathogenesis is through excessive synaptic pruning, a hypothesis with origins in older anatomical observations (Feinberg, 1982; Feinberg et al., 1967; Huttenlocher, 1979; Johnson and Stevens, 2018). One of the most consistent and robust neuro-imaging observations in schizophrenia has been of reduced brain volume associated with regional gray matter loss (Cannon et al., 2015; Haijma et al., 2013; Hu et al., 2015; Schmidt and Mirnics, 2015). Though several mechanisms have been posited (Harrison and Weinberger, 2005), many studies suggest a contribution from reduced synaptic content: meta-analysis of case-control post-mortem tissue studies found reductions in the presynaptic marker, synaptophysin, in hippocampus, frontal, and cingulate cortex (Osimo et al., 2018); individual and meta-analyses of histological studies have found reduced postsynaptic elements in layer III of prefrontal and temporal cortices (Glantz and Lewis, 2000; Sweet et al., 2009; van Berlekom et al., 2020); and a recent PET imaging study identified reduced levels of the synaptic vesicle protein SV2A in patient frontal and anterior cingulate cortex (Onwordi et al. 2020). Reduced synaptic content could be a consequence of insufficient synaptogenesis during early brain development or excessive synapse elimination (e.g. synaptic pruning) in later brain development. Developmental over-pruning was hypothesized early (Keshavan et al., 1994) as an interpretation, in part, of imaging studies showing enhanced neuropil contraction (increased neuropil catabolism) at first episode of psychosis and has been more recently supported by the observation of an increased rate of cortical thinning during the transition to psychosis in clinically high-risk adolescents (Cannon et al., 2015).

Outside the brain, activity of the complement cascade is a function not only of concentration of complement components, but also of the actions of complement inhibitory proteins, such as Decay Accelerating Factor (DAF), membrane cofactor protein (MCP), Factor H (FH), and C4b-Binding Protein (C4BP), which strongly influence the location and timing of complement deposition. For example, though complement is present throughout the blood, the presence of membrane-bound complement inhibitors on blood cells and absence of these inhibitors on microbes enables complement to specifically activate and deposit on microbial surfaces (Medzhitov and Janeway, 2002). If blood cells lose complement inhibitors then endogenous complement is activated and deposits on the blood cell surface, as is the case when somatic mutations lead to loss of DAF and CD59 in paroxysmal nocturnal hemoglobinuria (Brodsky, 2014). The complement inhibitors described above are predominantly expressed outside of the nervous system, and the landscape of complement inhibition in nervous tissue is unknown. CSMD1 may be well-situated to regulate brain-specific functions of the complement cascade, as *CSMD1* transcript is highly enriched in brain (The Genotype-Tissue Expression [GTEx] Portal; Distler et al., 2012; Escudero-Esparza et al., 2016; Escudero-Esparza et al., 2013; Steen et al., 2013).

Here we show, using both mouse tissue and human cell experiments, that endogenous CSMD1 acts as a complement inhibitor in neural tissues. We demonstrate that CSMD1 is selectively expressed in the CNS and predominantly by neurons, localizes to synapses, inhibits complement deposition, affects the development of a circuit in which synaptic pruning is regulated by complement and microglia, and alters the degree to which microglia consume synaptic components. Combined, these data show that loss of Csmd1 parallels the phenotype of C4 overexpression (Comer et al., 2020), support a model in which Csmd1 opposes complement cascade function in nervous tissues, and establish *CSMD1* as a second complement-related locus associated with schizophrenia.

## Results

### Loss of CSMD1 leads to enhanced deposition of complement on human neurons

To investigate whether endogenous human CSMD1 can act as a complement inhibitor in human neural cells, we used a human stem cell line genetically edited to knock out *CSMD1* (Hazelbaker et al., 2017). Stem cell-derived glutamatergic cortical-like neurons are one of the more well-characterized human neural cellular models, but it was unknown whether CSMD1 is expressed by glutamatergic cortical neurons in the human brain. Single molecule fluorescent *in situ* hybridization (smFISH) on *post mortem* human frontal cortex revealed that *CSMD1* mRNA was expressed broadly across cortical layers (Fig. 1a), and present in glutamatergic (expressing *Slc17a6*, which encodes vGluT2) neurons (Fig. 1b). We therefore differentiated human embryonic stem cells into glutamatergic cortical neurons (Nehme et al., 2018).

**Figure 1.**
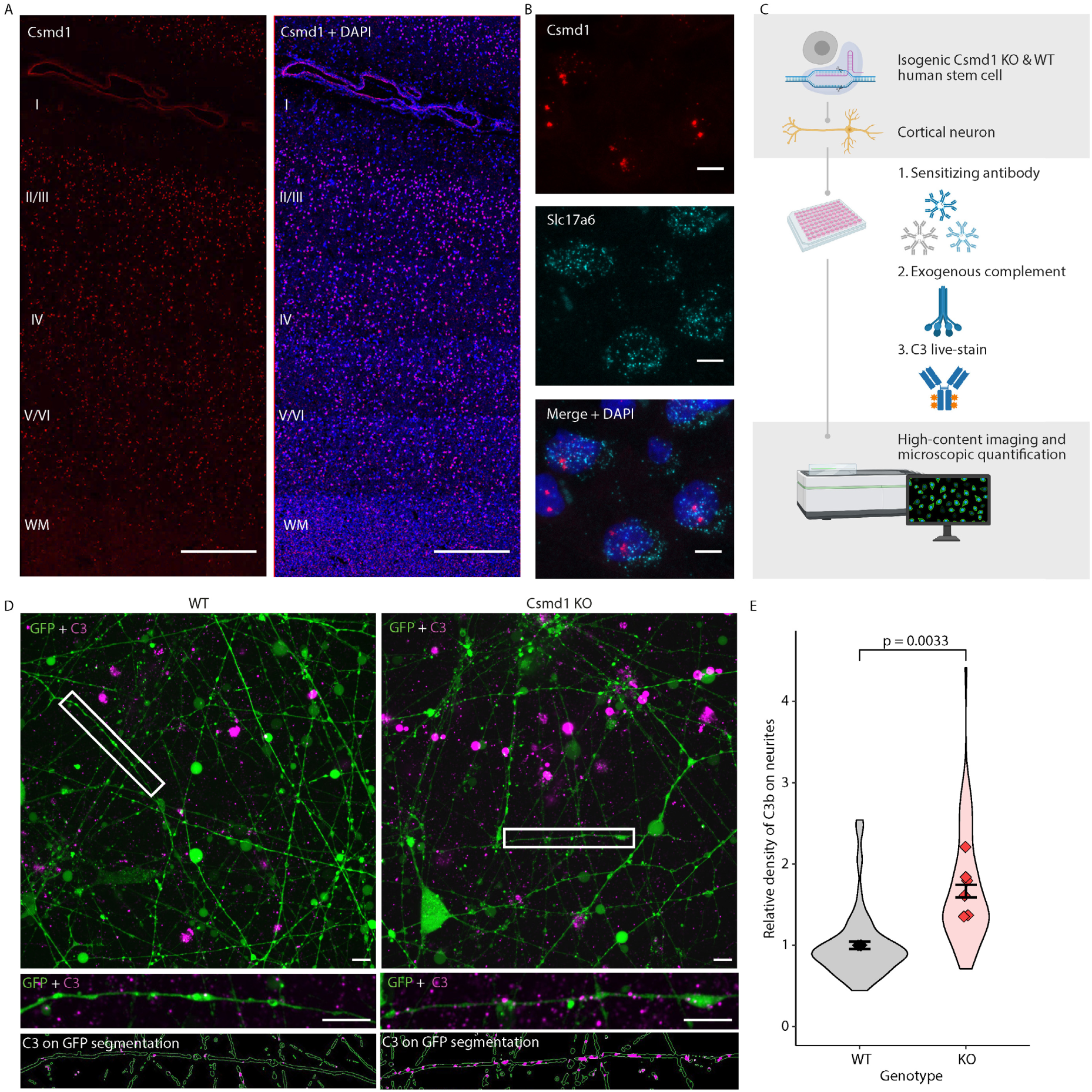
Human neurons lacking CSMD1 are more vulnerable to complement deposition. A) smFISH reveals *CSMD1* mRNA (red) expressed throughout grey matter layers of the *post-mortem* human cortex. White matter (WM) and approximate locations of cortical layers I-VI labeled in *CSMD1* alone image (left) and *CSMD1*+DAPI (DAPI = blue, on right). Scale bar is 500um and image is composed of tiled confocal z-stacks. B) Higher magnification images of smFISH (scale bar = 10um) show CSMD1 mRNA (red) is expressed in glutamatergic neurons (expressing *SLC17A6*, teal) in the human cortex. Image is a maximum intensity projection of a confocal z-stack. Merge includes DAPI (blue) C) A schematic of the complement deposition assay; see methods for details; created with biorender.com. D) Representative 63x maximum intensity projections of parental isogenic wild-type (WT) and CSMD1 knock-out (KO) human neurons (green) with live-stain for deposited C3b (magenta) with insets below of higher magnification neurites revealing C3b punctate deposition; scale bar in both overview and inset = 10um. E) CSMD1 KO neurons had more C3b deposition on their neurites compared to isogenic WT neurons; only C3b signal within a mask of GFP-positive processes was quantified and deposition was calculated as the sum of C3b intensity on GFP neurites, divided by neurite area. Each data point represents the average of all fields of view in an individual well, normalized to the average of WT wells on the respective plate. N = 79 wells WT and 74 wells KO pooled across 6 plates; p < 10^-11^ by Two-Tailed T-test. Error bars SEM. E) CSMD1 KO neurons had more C3b deposition on their neurites compared to isogenic WT neurons. Only C3b signal within a mask of GFP-positive processes was quantified and deposition was calculated as the sum of C3b intensity on GFP neurites, divided by neurite area. Data are presented as points representing plate means (N = 6) super-imposed on a violin plot of the distribution of individual wells (N = 79 WT and 74 KO). Well values are normalized to the mean of the WT wells within each plate. Two-Tailed T-tests: p= 0.0033 (plates) and p < 10^-11^ (wells). Error bars represent the mean ± standard error for all wells. See figure supplement 1e,f legend for additional statistical details.

To investigate whether loss of CSMD1 makes human neurons more vulnerable to complement, we challenged GFP-expressing differentiated neurons with a novel high-throughput complement-deposition assay. The assay is composed of three parts (Fig. 1c): 1) we catalyzed immune-complex formation on the surface of cells by applying a rabbit polyclonal antibody cocktail that broadly recognizes human cell-surface proteins; 2) we uniformly exposed neurons to normal human serum (NHS) containing all complement components; 3) we stained the live neurons for surface-deposited C3b. We then quantified C3b deposition onto neurites, restricting analysis to GFP-positive processes to enrich for healthy cells, as GFP signal is lost when cells become necrotic or apoptotic (Strebel et. al.; 2001; Arresate and Finkbeiner 2005). All branches of the complement cascade (the classical, alternative, and lectin pathways) converge on the activating cleavage of complement component C3 into C3a and C3b, the latter of which then deposits covalently on its target surfaces where it is recognized by microglial receptors to trigger engulfment in the brain. Thus, C3b deposition provides a robust readout of complement cascade activation.

We established that signal from the polyclonal anti-C3 antibody, which recognizes various forms of C3, showed a dose-dependent increase with concentration of complement-containing NHS added, and required complement activation, which is blocked by chelating requisite metal ions with EDTA – supporting an interpretation of this signal as reflecting cleaved, deposited C3b fragments (Fig. 1 – figure supplement 1a,b,c) (Escudero-Esparza et al., 2013). We found that human neurons lacking CSMD1 showed increased C3b deposition along their neurites compared to isogenic (i.e. “wild-type”) control neurons (Fig. 1d, e). Neurite density did not differ between genotypes (Fig. 1 – figure supplement 1d). Moreover the increased C3b deposition on neurons lacking CSMD1 was independent of the effect of density of neurites within each field (Fig. 1 – figure supplement 1e) or plate-to-plate variation (Fig. 1 – figure supplement 1f), supporting a model in which human CSMD1 expression on neurites inhibits complement cascade activity and may regulate complement deposition in the brain.

### Loss of Csmd1 enhances synaptic complement deposition and disrupts complement-dependent circuit development *in vivo*

Spurred by our observation that loss of CSMD1 rendered human neurons more vulnerable to complement deposition *in vitro*, we next sought to test the effect of loss of Csmd1 on the complement cascade *in vivo* in the mouse brain. Csmd1 protein is dramatically enriched in adult mouse brain lysates compared to other tissue lysates (Fig. 2A; Fig. 2 – figure supplement 1a) and is present broadly in the forebrain including subcortical structures like the thalamus and striatum (Fig. 2B,C; Fig. 2 – figure supplement 1b); this protein distribution is consistent with previous reports on mRNA distribution (Distler et al., 2012; Steen et al., 2013). We more closely examined the mouse retinogeniculate circuit (Fig. 2D), as developmental synaptic refinement within the dorsal lateral geniculate nucleus (dLGN) of the visual thalamus is well described (Corriveau et al., 1998; Guido, 2008; Hooks and Chen, 2006; Lee et al., 2014; Shatz, 1994; Stevens et al., 2007), and complement-mediated synaptic pruning contributes to this process (Stevens et al., 2007; Schafer et al., 2012; Lehrman et al., 2018). We found that 1) *Csmd1* mRNA is expressed in the retina, including by purified retinal ganglion cells (RGCs), and in the lateral geniculate nucleus (LGN) of the thalamus (Fig. 2E); and 2) that Csmd1 protein is abundant in the LGN by immunostaining (Fig. 2F).

**Figure 2.**
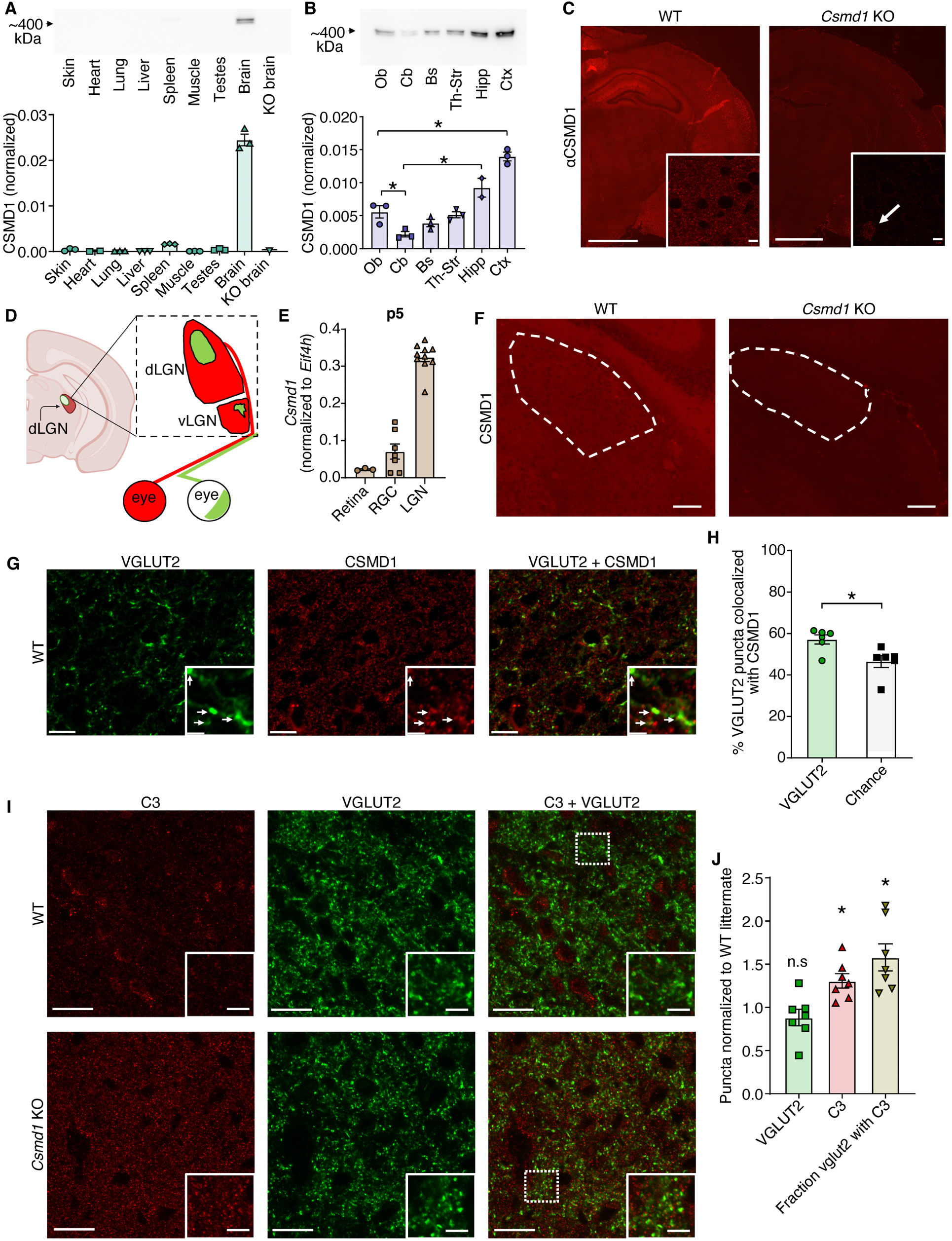
Loss of Csmd1 increases synaptic C3b deposition *in vivo* in a complement-dependent neural circuit. A-F, Csmd1 protein is highly enriched in the brain, where it is present broadly, including the Lateral Geniculate Nucleus (LGN) of the thalamus in the complement-dependent retinogeniculate circuit. A) Quantitative western blotting of total protein lysates from a panel of normal mouse tissues showed that Csmd1 is enriched in the brain; n=3 male p60 WT mice each tissue (n=2 for heart); brain enrichment is statistically significant with p < 1 × 10^-11^ by a post-hoc Tukey test after one-way ANOVA. Csmd1 was normalized to total protein stain (Bio-Rad stain-free gels) as shown in Figure 2 – figure supplement 1a. B) Quantitative western blotting of total protein lysates from p60 male mouse brain regions showed Csmd1 is present broadly in the brain; ob – olfactory bulb, cb – cerebellum, bs – brain stem, th-st – thalamus plus striatum, hipp – hippocampus, ctx – cortex, n = 3 animals (n=2 for hippocampus); * p < 0.05 post-hoc Tukey test after one-way ANOVA. Values are normalized to total protein (Bio-Rad stain-free technology) as depicted in Figure 2 – Figure Supplement 1B. C) IHC with a custom anti-Csmd1 antibody (Lund Custom made by Agrisera, Sweden) to an N-terminal sushi domain in Csmd1 showed broad staining of the neuropil in WT and non-specific nuclear staining in both WT and KO at p10; overview image is a mosaic of tiled images with scale bar = 5 mm. Inset shows 63x confocal images of layer II/III of somatosensory cortex; scale bar = 5 μm. D) Schematic of the mouse retinogeniculate system – the synapses from retinal ganglion cells (RGCs) in the eye to the lateral geniculate nucleus (LGN) in the thalamus; projections from each eye (red-contra, green-ipsi) compete for territory in the dLGN during development in a process dependent in part on complement mediated synaptic pruning by microglia, such that by p10, complement deposition has decreased and there is a modest area-overlap between the eye-specific territories. Created in part with biorender.com. E) *Csmd1* expression is detected by ddRT-PCR in the developing visual system at p5, an age of dynamic synaptic refinement. F) Csmd1 protein localizes to the neuropil of the dLGN (outlined) of a p10 WT mouse and is not seen in the dLGN of a KO mouse. Scale bar = 100 μm; these images are regions of interest from the coronal slices depicted in Fig 2C. G) At p10, Csmd1 puncta colocalized with vGluT2 puncta greater than would be expected by chance (represented by the colocalization when one channel was rotated by 90 degrees), scale bar = 10 µm overview and 2 µm inset, quantified in H, unpaired two-tailed t-test * p<0.05. I) There is an upregulation of C3b deposition in the dLGN at p10, when complement deposition is normally low and refinement of eye-specific territories is slowing. Representative images are single z-planes from WT and KO littermates; C3b (red), and retinogeniculate terminals (vGluT2, green), Scale bar = 20 µm in overview and 5um in inset. J) Quantification of C3b deposition reveals a greater number of C3b immunoreactive puncta and a greater fraction of vGluT2+ terminals colocalizing with C3b in the Csmd1 KO animals compared to their littermate pair; n = 7 WT and n= 7 KO littermates; quantification is shown as the Csmd1 KO level normalized to the WT littermate of each pair, where puncta counts for each animal (each “n”) are calculated as the per-FOV average across all FOVs for that particular animal; *p<0.05, one-way ANOVA with Dunnet’s multiple comparisons test; error bars are SEM. Brightness of representative images in C, F and I adjusted equally for display.

Complement deposition is observed at RGC synapses in the dLGN and contributes to developmental refinement and pruning of supernumerary retinal inputs during the first postnatal weeks (Stevens 2007; Schafer 2012). Anatomically, this developmental refinement contributes to “eye segregation”, in which initially overlapping inputs from the ipsilateral and contralateral eyes segregate into eye-specific territories in the dLGN – a central patch of ipsilateral inputs surrounded by contralateral inputs (Fig. 2D). Loss of complement components C1q, C3, or C4 results in reduced synaptic refinement, excess retinogeniculate synapses, and less eye-specific segregation (Stevens et al., 2007; Shafer et al., 2012; Sekar et al., 2016). We therefore assessed Csmd1 synaptic localization and C3b deposition in the dLGN at P10, when complement deposition is slowing in this brain region (Schafer et al., 2012), to test the hypothesis that loss of Csmd1 would result in greater complement deposition on retinogeniculate synapses. Csmd1 immunoreactive puncta in the early postnatal dLGN preferentially colocalized with vGluT2, a marker of glutamatergic RGC presynaptic terminals (Fig. 2G, H; Figure 2 – Fig. Supplement 2). Thus, Csmd1 is appropriately localized to regulate complement deposition at retinogeniculate synapses during the period of complement-dependent synaptic pruning and eye segregation. Furthermore, in concordance with the enhanced C3b deposition observed on *CSMD1* KO human neurons, immunostaining for C3 revealed an increase in the total number of puncta in the dLGN of Csmd1 KO mice compared to their WT littermates, and an increase in the fraction of vGluT2-positive retinogeniculate synaptic terminals colocalized with C3 (Fig. 2I,J).

To determine if loss of Csmd1 and increased synaptic complement deposition affects the refinement of eye-specific inputs in retinogeniculate circuit, we next examined eye-specific segregation of retinogeniculate synapses at the same age. *Csmd1* KO animals exhibited less binocular overlap than WT littermate controls (Fig. 3A, B, Fig. 3 – figure supplement 1c), consistent with increased synaptic refinement. RGC loss is unlikely to account for this phenotype (Blank et al., 2011), as we found no difference in RGC numbers in retinal whole mounts of WT vs KO animals (Fig. 3 – figure supplement 1a, b) and no genotype-specific difference in cross-sectional area of the contra- or ipsilateral dLGN (Fig. 3 – figure supplement 1d).

**Figure 3.**
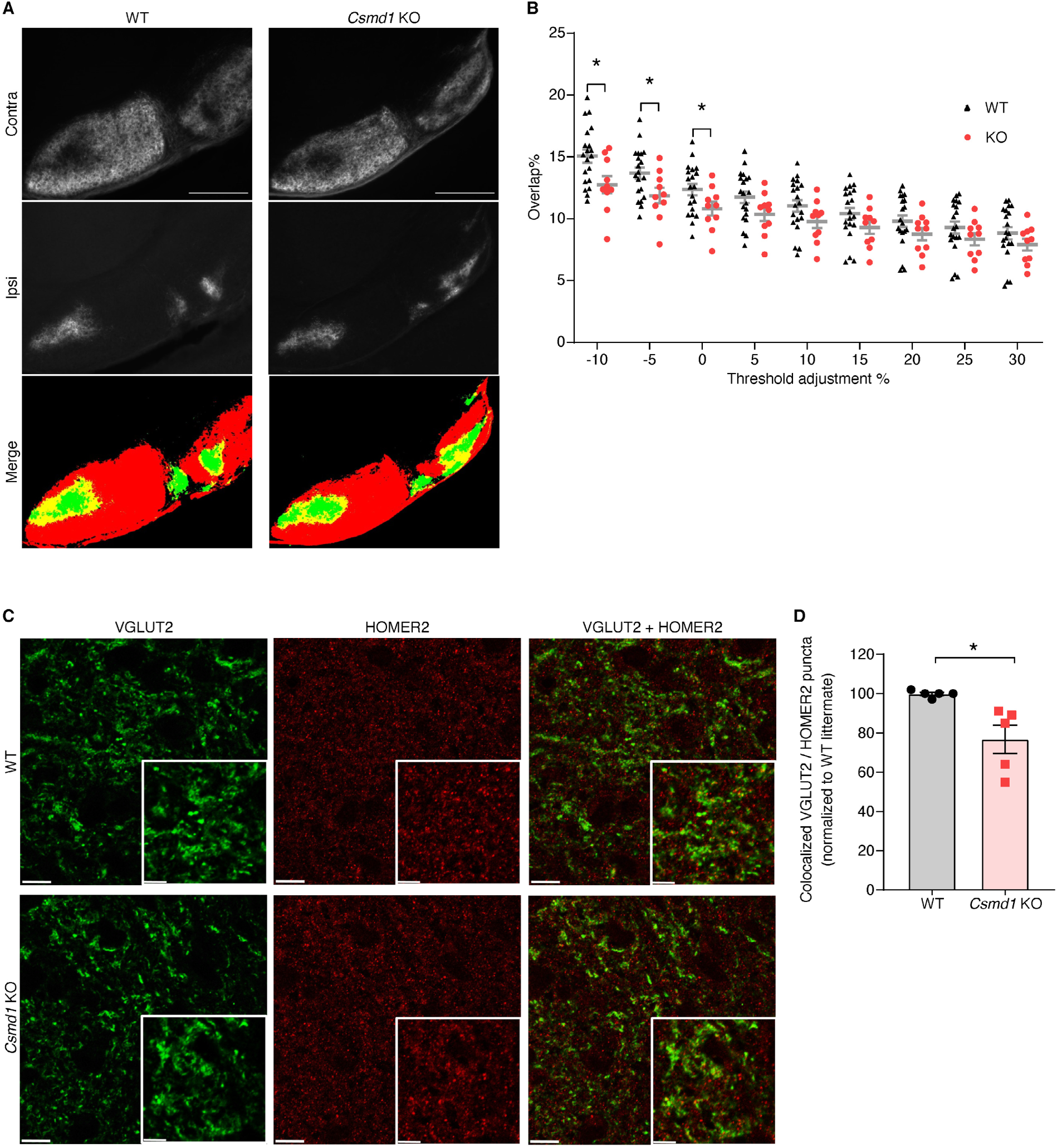
Loss of Csmd1 disrupts complement-dependent circuit development *in vivo*. A)-B) When complement-dependent segregation of retinal inputs to dLGN was assayed by labeling inputs from one retina in Alexa-488 and the other in Alexa-594, via cholera toxin beta subunit (CTB) injections to the retina and quantifying area of overlap in the dLGN at p10, we found that Csmd1 KO animals had less overlap of ipsi- and contralateral retinal territories in the dLGN than did WT; A) Representative images of contralateral inputs, ipsilateral inputs, and a merge of the binary-thresholded images of WT and KO animals. B) Quantification of overlap % (overlap area / total dLGN area) at 9 different binarization thresholds reveals a threshold-dependent effect of genotype where Csmd1 KO animals exhibited less overlap at lower thresholds. The baseline threshold stringency (0% adjustment) is subjectively determined by the individual analyst to separate background from fluorescent signal of fibers, and then the threshold adjusted by the percentage indicated on the x-axis; Because the curve of “overlap as a function of threshold-adjustment” risks artifactually flattening at higher stringencies in low-overlap phenotypes (e.g. if increasing the threshold stringency results in local “gaps” between parts of binary areas, further increasing threshold-stringency could not further decrease overlap in that local area), adjustments to make the baseline threshold more lenient (negative % adjustments, i.e. −10, −5%) were included as well as the set of adjustments to make the baseline threshold more stringent (positive % adjustment, i.e. +5, 10, 15, 20, 25, 30%) conventionally used to study phenotypes of high-overlap (e.g. in Stevens et al., 2007). * p < 0.05, unpaired two-tailed t-test at each threshold; error bars are SEM. By convention, each n is defined as an individual LGN with n=20 LGNs WT and 10 LGNs KO (animals WT=10, KO=5). C) Quantification of retinogeniculate synapses (colocalized vGluT2/Homer2 puncta) revealed that *Csmd1* KO mice have fewer of these synapses compared to their littermate controls; scale bars in overview are 10 μm and in inset, 5 μm. D) colocalized vGluTT2/Homer2 puncta quantification normalized to the WT(s) of littermate pairs; n=5 littermate pairs, p<0.05, unpaired two-tailed T-test; puncta counts for each animal (each “n”) are calculated as the per-FOV average across all FOVs for that particular animal.

Finally, consistent with a model in which CSMD1 influences synaptic density, *Csmd1* KO mice had a reduction in retinogeniculate synapse number, detected by immunostaining the P10 dLGN for puncta co-labeled with Homer2 and vGluT2. (Fig 3C, D; Figure 3 - Figure Supplement 2 A-C). The eye-segregation and retinogeniculate-synapse phenotypes seen in the absence of CSMD1 are similar to those seen in the absence of CD47 (Lehrman et al., 2018), an antiphagocytic ‘don’t eat me’ signal whose loss was previously shown to lead to enhanced synaptic pruning. The phenotype of the *Csmd1* KOs, moreover, is the opposite of models in which there is reduced complement-mediated synaptic pruning (KO of *C1q*, *C4*, or *C3*, or of the phagocytosis-enabling complement receptor, CR3, from microglia) (Schafer et al., 2012; Sekar et al., 2016; Stevens et al., 2007). The observations that loss of CSMD1 resulted in increased complement deposition, increased synaptic refinement (eye-specific segregation), and fewer retinogeniculate synapses, suggest increased complement-mediated synaptic pruning by microglia.

### *Csmd1* is expressed by a majority of cortical neurons and is present at synapses

Reports of synapse loss in cortical brain regions in people with schizophrenia (Glantz and Lewis, 2000; Sweet et al., 2009; Osimo et al. 2019; Onwordi et a., 2020) prompted us to further characterize the distribution of Csmd1 outside the retinogeniculate circuit. To investigate the dominant cell-types expressing *Csmd1* in a cortical region, we quantified *Csmd1* expression in nearly 1 million cells across three male p60 mouse frontal cortices using a high-throughput single molecule fluorescent in-situ hybridization (smFISH) platform (Fig 4a-f). We found that approximately 4 out of 5 *Csmd1*-expressing cells in frontal cortex were either GABAergic or glutamatergic neurons and that glutamatergic neurons comprised the majority of *Csmd1* positive cells (Fig. 4e,f); approximately one third of total cells analyzed (∼285,000) expressed *Csmd1*. We found that ∼80% of GABAergic neurons and ∼60% of glutamatergic neurons expressed *Csmd1*, suggesting that *Csmd1* is expressed by a majority of cells within these neuronal types but also that the expression may be subclass or cell-state specific. These results further show that the expression of *CSMD1* by human glutamatergic neurons of the frontal cortex (Figure 1) is conserved in mice.

**Figure 4.**
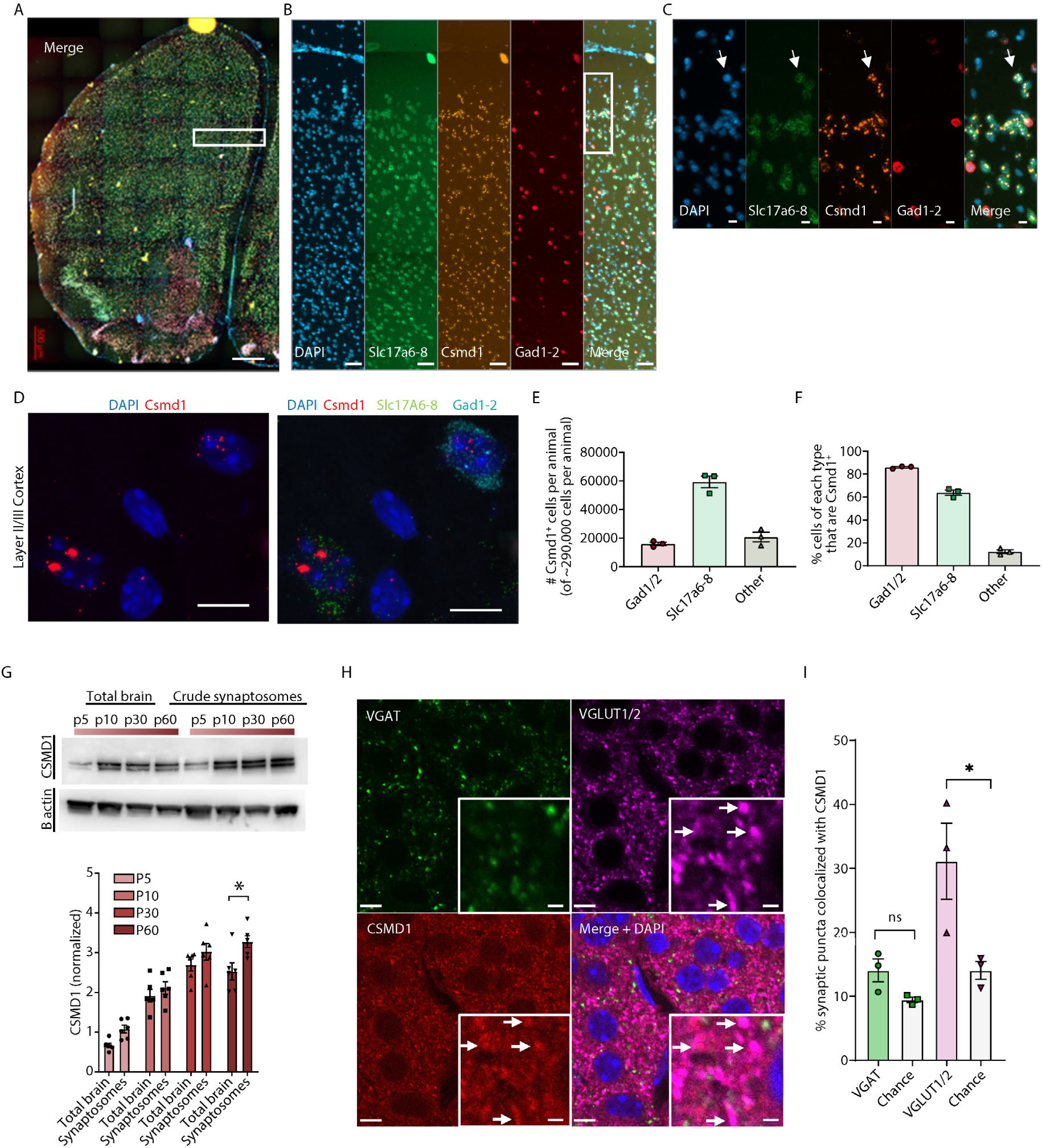
*Csmd1* is expressed by the majority of neurons in cortex, and its protein localizes to synapses. A-C, E-F) smFISH of p60 mouse frontal cortex showed *Csmd1* mRNA (orange) expression in glutamatergic (*Slc17a6-8*, green) and GABAergic (*Gad1-2*, red) neurons, and in other cell populations (marked by only DAPI, blue); n = 3 animals, ∼290,000 cells counted per animal with a custom CellProfiler pipeline. Images are tiled 20x fields of view. A) Overview of coronal slice from one hemisphere; scale bar = 500um. B) Expression was seen throughout the cortical layers scale bar = 50 μm. C) Inset near surface of cortex; *Csmd1* mRNA can be seen as 2-3 large nuclear clusters of puncta and scattered cytoplasmic puncta. A *Csmd1*-positive cell that is a probable non-neuronal cell can be seen at the top of the inset. D) 100x resolution maximum intensity projection of confocal z-stack of smFISH visualizing individual puncta of *Csmd1* in glutamatergic (Slc17a6-8+) and GABAergic (Gad1-2+) neurons in layer II/III of frontal cortex of p60 mouse brain (*Csmd1*, red; *Slc17a6*-*8*, green; *Gad1-2*, teal; DAPI, blue). E) The cell-type breakdown of *Csmd1* expression showed that it was expressed predominantly by neurons. F) Quantification of the fraction of GABAergic neurons (∼80%) and glutamatergic neurons (60%) and other cells that express *Csmd1* revealed a heterogeneity within each cell class. G) Quantitative western blot of total brain homogenates and crude synaptosomal fractions (P2) showed that Csmd1 was present in synaptosomes and that its concentration increased with age; effect of age (p < 0.05) and biochemical fraction (p < 0.05) with two-way ANOVA. A post-hoc Sidak test for multiple comparisons showed that Csmd1 is enriched in synaptosomes compared to total brain at p60 (*p < 0.05); n= 6 animals each age. Csmd1 is normalized to B-actin. H,I) Csmd1 colocalizes with vGluT1/2 excitatory synaptic puncta more than expected by chance (as simulated by rotating one image channel by 90 degrees) in p10 cortex; p < 0.05 two-way ANOVA with Sidak multiple comparisons test, N=3 animals, Scale bar = 5 μm overview and 1μm inset, vGat (green), pooled vGluT1 and vGluT2 (magenta), Csmd1 (Red), DAPI (Blue); arrows show several vGluT/Csmd1 colocalized puncta. Image brightness was adjusted for visualization only. The frontal cortex smFISH dataset used in A-F was also used to estimate the E/I (excitatory/inhibitory) neuron ratio as described in Saunders et al., 2018 (Figure S4 G, H.)

We next investigated whether CSMD1 might be found at synapses in the cortex. Staining of coronal sections of the p10 mouse brain revealed puncta in neuropil and in synapse-dense regions (such as the synaptic layers of the hippocampus) and low signal in white matter tracts (Fig 2C, Fig 2 – Fig Supplement 1D). CSMD1 protein was increasingly abundant in synaptosomes – a brain cellular fraction enriched for synaptic components (Luquet et al., 2017) – over postnatal development, and became significantly enriched in the synaptosome fraction compared to total brain as the animals transitioned to sexually-mature adulthood (p60) (Fig. 4g, Fig. 4. Fig. Supplement 1). Finally, CSMD1 immunoreactive puncta significantly colocalized with markers for glutamatergic synapses (vGLUT1/2+) in cortex at P10 (Figure 4H, I), demonstrating that CSMD1 is present at a subset of synapses even at this early developmental stage. Combined, these results suggest that CSMD1 is expressed by a majority of excitatory and inhibitory cortical neurons, is broadly present at cortical synapses, and exhibits neuronal, synaptic, and developmental heterogeneity, making it well-positioned to regulate cortical synaptic pruning and circuit refinement.

### Loss of *Csmd1* enhances engulfment of synaptic elements by microglia *ex vivo*

Complement components are ‘eat me’ signals that mediate the engulfment of synapses by microglia during development; genetic ablation of complement components reduces the amount of phagocytosed synaptic material inside microglia. To examine whether loss of *Csmd1* changes the engulfment of synapses by microglia, we used an assay in which fluorescently labeled synaptosomes are “fed” *ex vivo* to cultured microglia (Fig 5a; Fig. 5 - Figure Supplement 1 a, e). This type of *ex vivo* engulfment assay had previously been useful for detecting enhanced synaptic engulfment due to loss of the phagocytic inhibitor CD47 (Lehrman et al., 2018). When synaptosomes from whole forebrain were “fed” to WT mouse microglia, the microglia engulfed more synaptosomes purified from CSMD1 KO mice compared to synaptosomes from WT littermates (Fig 5 b, c; Fig. 5 - Figure Supplement 1 b, c, d). Moreover, the rate of engulfment of synaptosomes from *Csmd1* KO animals was faster (Fig. 5d). Combined, these data show that loss of *Csmd1* enhances *ex vivo* engulfment of synaptic elements by WT cultured microglia, which suggests that Csmd1 may regulate engulfment of synapses by microglia.

**Figure 5.**
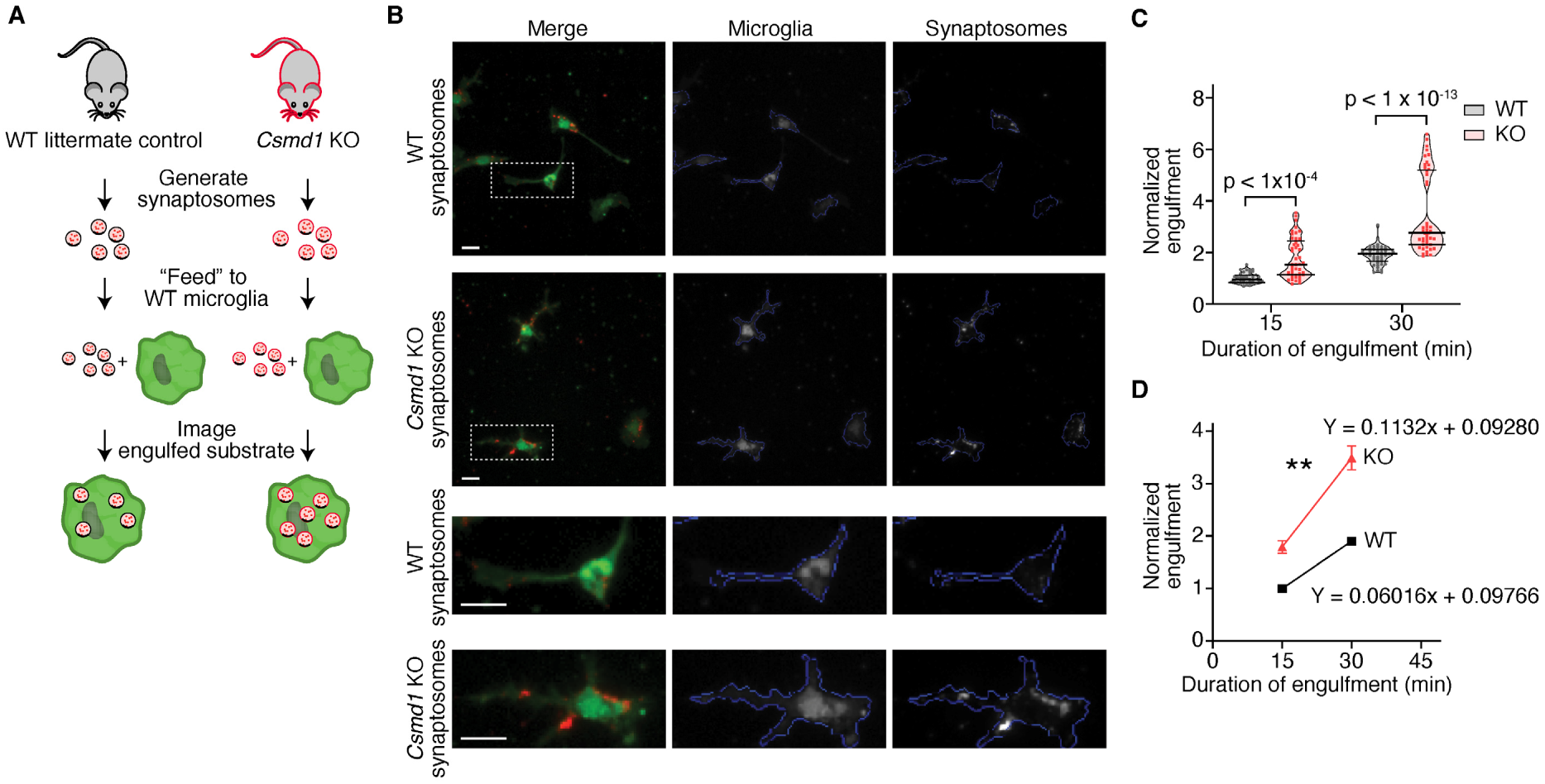
Loss of *Csmd1* enhances the *ex vivo* engulfment of synaptic components by cultured WT microglia. A) Schematic of *ex vivo* synaptosome engulfment assay where crude synaptosomes were isolated from WT and *Csmd1* KO sex-matched littermates, labeled with fluorophore, fed to cultured WT microglia, and the fluorescent signal within microglia quantified as a proxy of engulfed synaptic material. A high-content imaging pipeline was used to delineate individual cells and synaptosome fluorescence was measured per cell. B) Representative images of WT microglia fed either WT or *Csmd1* KO synaptosomes. Scale = 10 μm C) Microglia engulf more synaptosome substrate from *Csmd1* KO animals compared to WT at both 15min and 30min after addition of synaptosomes. Synaptosome engulfment is the fluorescence intensity within microglia for each well, normalized to the average of the WT wells at 15 min on each plate. Each data point represents the average of 64 FOVs from a single well (n = 45 wells each genotype each timepoint) pooled from 3 separate experiments; see Figure 5 – figure supplement 1 for individual experiments. P < 1 × 10^-4^ (15 min) and P < 1 × 10^-13^ (30 min), two-way ANOVA with Sidak’s multiple comparisons test. D) The rate of engulfment (represented by the slope of a linear regression between 15 min and 30 min) of *Csmd1* KO synaptosomes is greater than of WT synaptosomes, ** p < 0.01. Goodness of fit of linear model: R^2 = 0.72 (WT) and 0.33 (KO).

## Discussion

Though *CSMD1* was one of the earliest genes implicated in GWAS studies of schizophrenia, its functions have remained largely unknown. Previous studies have shown roles for *CSMD1* in the development of sperm and mammary ducts (Lee et al., 2019) and as a putative tumor suppressor in cancer (Blom, 2017; Escudero-Esparza et al., 2016; Kamal et al., 2010; Ma et al., 2009; Shull et al., 2013; Zhang and Song, 2014), and have demonstrated complement regulatory potential *in vitro* (Escudero-Esparza et al., 2013). Previous studies of *Csmd1* KO mice have been restricted to general behavioral assays, the interpretation of which was confounded by the mice having a mixed genetic background (Distler et al., 2012; Drgonova et al., 2015; Steen et al., 2013). Here, we presented evidence that endogenous CSMD1 opposes complement activities in neural tissues. Loss of *CSMD1* rendered human stem-cell derived neurons more vulnerable to activated complement leading to complement deposition on neurites. Experiments revealed regional and cell-type heterogeneity of *Csmd1* expression in the mouse brain. Loss of *Csmd1* increased complement deposition, decreased synapse number, and abrogated the development of a circuit that undergoes complement-dependent synaptic pruning. More broadly, Csmd1 protein was expressed across brain regions, present at diverse synapses, and affected the extent of synaptic engulfment by microglia in culture.

### A mechanism to enable differential vulnerability to complement-dependent processes

The grey matter loss and synaptic deficits described in schizophrenia exhibit regional, cellular, synaptic, and temporal specificity that remains under-explained. Prefrontal cortex, cingulate cortex, and hippocampus are most affected by grey matter loss, for example, although the region specificity is still debated (Chua et al., 2007; Haijma et al., 2013; Whitford et al., 2006). We found *Csmd1* to be differentially expressed by GABAergic and glutamatergic cortical neuron types, an observation reinforced by recent single cell RNA-sequencing data (Saunders et al., 2018); and that the level and synaptic localization of the protein changes with development. This raises the question of whether Csmd1 could affect which ages, regions, neurons, and synapses are more or less vulnerable to complement-dependent processes. As a recent paper found that inducing retinal ganglion cells to exogenously express Crry, a known complement inhibitor, protects not only the retinogeniculate synapses from complement deposition and microglial engulfment, but also neighboring synapses (though to a lesser degree) (Werneburg et al., 2020), non-cell autonomous effects could add another layer of complexity. Much of our mechanistic study focused on glutamatergic cells and synapses, and we found Csmd1 to be preferentially co-localized with glutamatergic synaptic markers in early postnatal mouse cortex. However, a recent proteomic characterization of cultured rat cortical neurons found Csmd1 was indeed part of the inhibitory (neuroligin-2 positive) synapse proteome (Loh et al., 2016), leaving open the possibility that Csmd1 could play similar or different roles at inhibitory synapses.

We showed that loss of *Csmd1* leads to more C3b deposition and enhanced synaptic refinement in the retinogeniculate system. Moreover, Csmd1 protein is present broadly elsewhere in the brain, including cortex and hippocampus, and cultured microglia engulfed more bulk forebrain synaptosomes in the absence of Csmd1, but whether Csmd1 may regulate synapse numbers or the complement system in these regions remains unknown. Though complement-mediated synaptic pruning is less established in these regions, it has recently been reported that exogenously increased complement levels can reduce synapse numbers, alter connectivity, and increase microglial engulfment in mouse frontal circuits (Comer et al., 2020).

### A broader landscape of neural complement regulation

We believe this work to be one of the first description of a neural complement inhibitor and might serve as a catalyst for investigations of other putative neural complement inhibitors. *CSMD1* has two paralogues, *CSMD2* and *CSMD3*, which each share ∼60% encoded protein sequence homology with *CSMD1* and are highly similar in their domain structure (differing marginally in the number of sushi domains); all *CSMD* transcripts appear to be enriched in brain (gtexportal.org). Knock-down of *CSMD2* has recently been shown to lead to decreased arborization and decreased spine density in cultured hippocampal neurons (Gutierrez et al., 2019, 2020), although the involvement of complement in this function remains untested. Within the *CSMD1* locus itself, moreover, there may be distinct functions of alternative isoforms that have been reported in public transcriptome databases, but have not been characterized. Both the human and mouse genetic models, here, disrupt an initial exon to create a loss of function model and eliminate full length protein (confirmed by western blotting, IHC, and ICC), but may not disrupt proteins generated from downstream alternative start sites should they exist. However, two lines of evidence suggest that if these alternative transcripts are translated, they are minor isoforms. First, it is known that the *Csmd1* KO mouse has residual expression of mRNA from exons downstream to the deletion/insertion in exon 1 (Steen et al., 2013) but we did not detect residual protein expression in western blot at a size predicted by any of these predicted alternative transcripts; as the western blot antibody recognizes an epitope in the transmembrane domain nearly at the C-terminus of the protein, it would theoretically detect isoforms if they were abundant. Second, a sparse, nuclear signal was observed in both WT and KO with a polyclonal antibody to Csmd1, but pre-absorption of the antibody for 5 days on KO brain lysate prior to immunostaining eliminated the nuclear signal but not the punctate neuropil signal, suggesting that the nuclear signal was due to an off-target antibody in the polyclonal mixture, not to residual expression of a *Csmd1* isoform (data not shown). These observations do not obviate a role for possible minority alternative Csmd1 isoforms, however.

Besides the CSMD1, 2, and 3 proteins, there are only a handful of other known proteins that contain both CUB and sushi domains, and these appear only in vertebrates. The only one of these with a known function is Sez-6, which was described to regulate the number and maturity of excitatory synapses and the extent of dendritic arborization in mouse neuronal culture and *in vivo* (Gunnersen et al., 2007; Gunnersen et al., 2009). Experiments in Sez-6 KO neuronal cultures found a reduction in spine maturity, i.e. spines with heads *vs*. without heads (Gunnersen et al., 2007). Three sushi-domain-containing proteins, Susd2, Sez-6 (as above), and Srpx2 have been shown to regulate excitatory synapse number in culture and/or *in vivo* in mouse (Gunnersen et al., 2007; Nadjar et al., 2015; Sia et al., 2013; Soteros et al., 2018); evidence for a connection with the complement cascade and pruning has recently been presented for Srpx2 (Cong et al., 2020) but has not been investigated for the other sushi-domain containing proteins. Among these, only *CSMD1* has been consistently associated with schizophrenia to date.

### Csmd1 and the etiology of schizophrenia

In the context of a pruning hypothesis of schizophrenia, our data that CSMD1 opposes complement activity suggest a model in which reduced expression or function of CSMD1 could increase schizophrenia risk through enhanced complement-mediated pruning. Precisely how risk alleles in intron 3 of *CSMD1* may change the gene’s regulation in a way relevant to schizophrenia is unknown. Such variation could in principle alter global, regional, cell-type, or developmental regulation of *CSMD1* expression; nuclear retention of transcript, which is common for long genes (Bakken et al., 2018; Tuffery-Giraud et al., 2017); or susceptibility to double-stranded DNA breaks known to occur at this locus (Wei et al., 2018; Wei et al., 2016). Yet the relationship between synaptic pruning and cognitive impairments could provide a clue to the role of *CSMD1* polymorphism in schizophrenia. Cognitive impairments are seen in several models of brain disease in which enhanced complement activity and microglial dependent synaptic pruning have been implicated, including Alzheimer’s disease, and West Nile virus-induced memory impairment (Hong et al., 2016; Schafer et al., 2016; Vasek et al., 2016). Cognitive impairments also present an important unmet clinical need in schizophrenia (Collins et al, 2011; Hyman, 2013) and the extent of grey matter loss is positively correlated with the duration of untreated psychosis and magnitude of functional impairment (Cahn et al., 2006; Malla et al., 2011). Although currently there has been only limited study of possible effects of CSMD1 on cognitive function, genetic variation in *CSMD1* has been reported to affect cognitive and executive function in healthy adult humans (Athanasiu et al., 2017; Koiliari et al., 2014), and several studies reported reductions in measures of cognitive functioning in individuals with the risk allele of the schizophrenia-associated common variant in *CSMD1*, although sample sizes were small (Donohoe et al., 2013; Rose et al., 2013).

The work presented here uncovers new biology of the schizophrenia risk gene, *CSMD1* and suggests that CSMD1 opposes the activity of complement in neural tissues. Though we have highlighted relevance to schizophrenia, complement, and synaptic pruning, CSMD1 may also have other neural functions, and is implicated in other disorders. Recent work has associated rare genetic variations in *CSMD1* with Parkinson’s disease, for example (Ruiz-Martínez et al., 2017). It will be important, consequently, to further expand biological understanding of CSMD1 and uncover the breadth of its functions in normal brain development and illness.

## Materials and Methods

### *Csmd1* KO Mice

The *Csmd1* KO mouse was originally created by Lexicon Pharmaceuticals through deletion of a 1 kb genomic sequence composed of part of exon 1 and intron 1 caused by insertion of a *LacZ/Neo* cassette and rederived by Taconic on a mixed B6N:129 background (Distler et al., 2012; Steen et al., 2013). The KO was then backcrossed over ∼2 years onto a pure C57bl6N-Taconic background to control for effects of the mixed background; SNP genotyping of breeders found them to be 98.17% to 99.37% C57BL/6N-Taconic (Taconic 1449 MD SNP Linkage Panel). Though this model has been used in several published studies, it is known that there is residual transcription from the *Csmd1* locus (Steen et al., 2013) so we confirmed that it has lost Csmd1 protein, as assayed by an antibody against an antigen in the transmembrane (TM) domain (Abcam); since the TM domain is encoded by the 3’ end of the mRNA in the region where previous studies detect residual transcript, the result increases the likelihood this mouse is a true KO at the protein level. Mice were group housed in Optimice cages and raised in a facility according to conditions recommended by AAALAC. All procedures were approved conducted in accordance with guidelines from the Boston Childrens Hospital Institutional Animal Care Use Committee, as well as guidelines from the National Institutes of Health.

### Mouse Genotyping

*Csmd1* KO mice were genotyped with a custom assay using the Phire Tissue-direct PCR kit (Thermo Fisher Scientific) according to the manufacturer’s “dilute and store” protocol. Briefly, single mouse toes were incubated with 50 ul of dilution buffer and 1.25 ul DNARelease additive for 2 min at 98°C to liberate DNA. A 2-step multi-plexed PCR protocol was followed using 1 ul of the above solution as input in a 20 ul total volume reaction. Reagent setup (left) and thermocycling (right) was as follows:

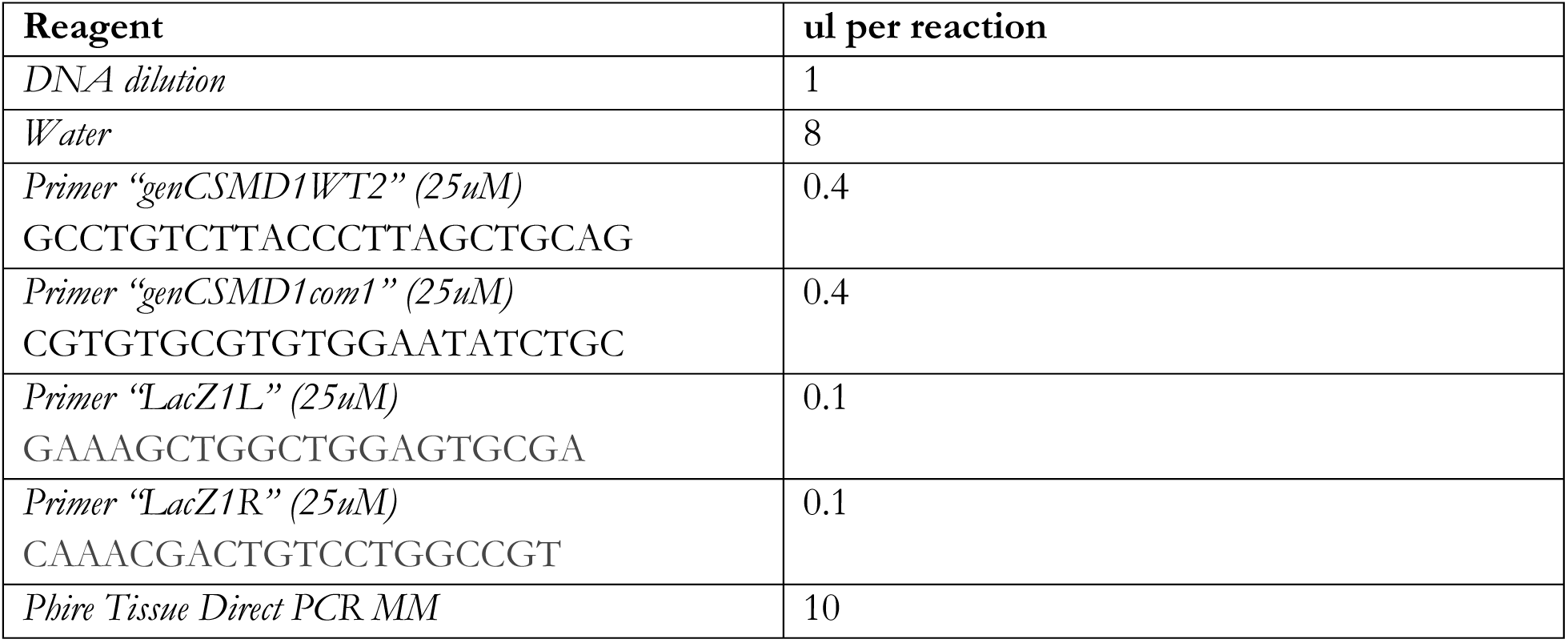

| Step | Temperature | Time |
| --- | --- | --- |
| 1 | 98°C | 5 min |
| 2 | 98°C | 5 sec |
| 3 | 72°C | 20 sec |
| 4 | Go to Step 2 | 39 cycles |
| 5 | 72°C | 1 min |
| 6 | 4°C | Forever |

The PCR product was visualized on 2% agarose gels with ethidium bromide in which the WT band is 122 bp and the KO band is 291 bp.

### Complement deposition on human derived neurons (Figure 1D-F; Figure 1 – Figure Supplement 1)

All embryonic stem cell work was performed according to an approved ESCRO protocol from Harvard University and the Broad Institute. *CSMD1* KO and isogenic WT human embryonic stem cell lines (Hazelbaker et al., 2017) were differentiated using viral doxycycline-inducible overexpression of NGN2 (Nehme et al., 2018), co-infected with eGFP expressing virus, and cultured in 96-well, glass bottom plates (Senseo Greinier) for 14 days *in vitro* in serum-free medium without feeder cells. At day 14, a complement deposition assay was performed, modified from (Escudero-Esparza et al., 2013) to apply to cultured ES-derived neurons. All solution changes were carried out with an Integra VIAFLO 96/384 with 96 channel head and tips, leaving just enough liquid behind at each step to keep cells submerged at all times to maximize cell health and adhesion. Cells were sensitized to the classical complement cascade through the generation of immune complexes on their surfaces by treating for 20 min at room temperature with 50 ug/ml custom rabbit pan cell-surface polyclonal antibody generated against Bjab/Jurkat/Thp-1 cells (Custom made by Agrisera, Sweden and generously provided by Anna Blom’s group, Lund, Sweden) in binding buffer (10 mM Hepes, 140 mM NaCl, 5 mM KCl, 1mM MgCl_2_, 2 mM CaCl_2_, 0.02% azide, pH 7.2). Cells were washed in either GVB++ or GVBE buffer (Complement Technology) and incubated for 30 min at 37°C with a source of exogenous complement: either 3% normal human serum (NHS) in either GVB++ (permissive of complement deposition) or GVBE (negative control – not permissive of complement deposition due to presence of EDTA) (Figure 1 D-F, Figure 1 – Figure Supplement 1 C, D) or a range of 0%-10% NHS in either GVB++ or GVBE (Figure 1 – Figure Supplement 1 A, B). Cells were washed 2x and live stained in warm neurobasal media with Dylight-650-conjugated (Abcam) rabbit anti C3 antibody (DAKO) 1:200 for 20 minutes at 37°C. Cells were washed in neurobasal media and fixed 7 minutes in 4% PFA, followed by several washes in PBS.

Plates were imaged on a Perkin Elmer Opera Phenix High Content Confocal system with 63x water immersion objective. Eight fields of view per well were captured as 6-plane z-stacks spanning 2.5 µm, and maximum intensity projections analyzed in the attached Harmony analysis software. Empty wells, or notably damaged wells were flagged from 10x overview images and removed from analysis; well values were obtained as the average of all FOVs in that well. C3b signal within a mask of GFP-positive processes was quantified and deposition was calculated as the sum of C3b intensity on GFP neurites, divided by neurite area.

NHS was obtained from consenting healthy adults according to the recommendations of the local ethical committee in Lund, Sweden (permit 2017/582). Blood samples from 11 adults was allowed to clot 30min at 25°C, centrifuged at 1500g for 20min at 4°C to separate the NHS, then NHS was pooled, aliquoted and stored at −80°C until immediately before use.

### Visualization of *Csmd1* with single molecule fluorescent in-situ hybridization (Figure 1 A,B; Figure 4 A-F)

For high-throughput characterization of *Csmd1* expression in mouse frontal cortex (see Figure 2 – Figure Supplement 1b for methods schematic), three P60 male mice were deeply anesthetized with isoflurane, cervically dislocated, decapitated and brains extracted. Brains were snap frozen on an aluminum foil boat floating in liquid nitrogen, and embedded side-by side in OCT (VWR). The brains were sectioned (14 μm/section) on a Leica cryostat and kept cold on dry ice while avoiding over-desiccation. Every 5^th^ section was processed for single molecule fluorescent in situ hybridization using the Advanced Cell System’s (ACD) branched DNA technology (RNAscope) according to the manufacturer’s protocol for fresh frozen tissue with multiplexed fluorescent amplification (ACD user manual numbers 320513 and 320293). Briefly, slides were post-fixed for 15 minutes in freshly prepared 4% PFA (Electron Microscopy Sciences) in PBS followed by alcohol dehydration. Sections were permeabilized with “pretreat IV” and then hybridized using probes to *Csmd1* (channel 1), pooled probes to *Slc17a6*, *Slc17a7*, and *Slc17a8* (all channel 2), and pooled probes to *Gad1* and *Gad2* (channel 2); see reagents table for target probe details. Tissue quality was qualitatively assessed as having robust and uniform signal on the RNAscope Triplex Positive control (containing probes for high, medium and low abundance transcripts), and minimal signal on Triplex negative control (DapB, a bacterial transcript) according to the manufacturer’s recommendation. “Amp B” was used to visualize the probes, putting channel 1 in red, channel 2 in green, and channel 3 in far red. After the procedure, slides were briefly counterstained with DAPI and cover slipped using ProLong Gold Antifade Mountant (ThermoFisher); note, the multiplexed fluorescent amplification is incompatible with glycerol-based mounting media like Vectashield (Vector Labs). Slides were scanned on a Zeiss Axioscan.Z1 with a 20x objective and quadruple bandpass filter set. For cell counting experiments, images were converted to TIFF via the Zeiss CZItoTiffBatchConverter (CZI to Tiff Converter Suite software by Zeiss) with downsampling (zoom factor x 4) to speed downstream processing. Frontal cortex was identified pseudo-manually by drawing regions of interest (ROI) in a custom CellProfiler (2.1.1) pipeline; any meninges and torn or folded regions in slices were excluded from ROIs. In a second custom CellProfiler pipeline, ROIs were imported as masks and cells positive for smFISH labeling of *Csmd1*, *Slc17a6-8*, and *Gad 1-2* were automatically identified and counted: briefly, the pipeline included illumination correction, smoothing of objects, thresholding, object indentation and splitting of clumped objects. Objects in different channels were counted as colocalized if the object centroid in one channel lay within a 2-pixel radius on either side of the centroid of the object in the other channel (i.e., a 5-pixel diameter circle). As quality control measures, object numbers per image were plotted and outliers manually inspected. For absolute numbers of cells identified between animals, the atlas region analyzed was trimmed to match the animal with the most anterior 1^st^ slice; for relative numbers (proportions of cells per animal), all slices passing quality control from each animal were included. This dataset was also used to determine the normal E/I ratio of mouse frontal cortex in (Saunders et al., 2018) as detailed in the Figure 2 legend. For high resolution image of mouse cortex (Figure 4 D), a maximum intensity projection of a z-stack of layer II/III of frontal cortex was imaged at 100x on a Leica SP8 with a white laser.

For smFISH in human tissue, the proteinase digest step of the RNAscope procedure was extended to 30min but was otherwise the same as above. Probes for human *CSMD1* and *SLC17A6* were used as detailed in reagent table. Images were captured as 63x confocal z-stacks on a Leica SP8; images were tiled to form a mosaic in fig 1A. Control human brain tissue imaged in Figure 1 was obtained from the McLean brain bank; demographics are as follows: Female, age 76, post-mortem interval 8.08. As there is much variability in the RNA quality of banked human tissue, this tissue sample was selected based on good signal with the RNA-scope triplex positive control.

Frozen human brain tissue was obtained from the Harvard Brain Tissue Resource Center (HBTRC; McLean Hospital), which acquires de-identified postmortem human brain tissue under approval from the Partners Human Research Committee and with permission from legal next-of-kin for the use of brain tissue for research. Federal regulation 45 CFR 46 and associated guidance indicates that the generation of data from de-identified postmortem specimens does not constitute human subjects research requiring institutional review board review. Postmortem tissue collection followed the provisions of the United States Uniform Anatomical Gift Act of 2006 described in the California Health and Safety Code section 7150 and other applicable state and federal laws and regulations. History of psychiatric or neurological disorders was ruled out by consensus diagnosis carried out by retrospective review of medical records and extensive questionnaires concerning social and medical history provided by family members. Several regions from each brain were examined by a neuropathologist. Tissue used for this study did not include subjects with evidence of gross and/or macroscopic brain changes, or clinical history, consistent with cerebrovascular accident or other neurological disorders. Subjects with Braak stages III or higher (modified Bielchowsky stain) were not included. None of the donors had significant history of substance dependence within 10 or more years of death, as further corroborated by negative toxicology reports.

ACD RNAscope Probe Table.

| Gene Symbol | Gene Name | Species | Accession | ACD Catalog Number | Channel | Probe Pairs | Probe Region Begin | Probe Region End |
| --- | --- | --- | --- | --- | --- | --- | --- | --- |
| *CSMD1* | CUB and Sushi multiple domains 1 | Homo sapiens | NM_033225.5 | 424631 | 1 | 25 | 255 | 1369 |
| *Csmd1* | CUB and Sushi multiple domains 1 | Mus musculus | NM_053171.2 | 444791 | 1 | 20 | 263 | 1489 |
| *3-plex positive control* | peptidylprolyl isomerase B, RNA polymerase II subunit A, ubiquitin C | Homo sapiens |  | 320861 | 3 |  |  |  |
| *3-plex negative control* | Bacterial Gene DapB | Any Species |  | 320871 | 3 |  |  |  |
| *3-plex positive control* | peptidylprolyl isomerase B, RNA polymerase II subunit A, ubiquitin C | Mus musculus |  | 320881 | 3 |  |  |  |
| *Gad1* | glutamate decarboxylase 1 | Mus musculus | NM_008077.4 | 400951-C3 | 3 | 15 | 62 | 3113 |
| *Gad2* | glutamic acid<br>decarboxylase<br>2 | Mus<br>musculus | NM_008078.2 | 439371-C3 | 3 | 20 | 552 | 1506 |
| *Slc17a6* | solute carrier<br>family 17<br>(sodium-<br>dependent<br>inorganic<br>phosphate<br>cotransporter),<br>member 6 | Mus<br>musculus | NM_080853.3 | 319171-C2 | 2 | 20 | 1986 | 2998 |
| *Slc17a6* | solute carrier<br>family 17<br>(sodium-<br>dependent<br>inorganic<br>phosphate<br>cotransporter),<br>member 6 | Homo<br>sapiens | NM_020346.2 | 415671-C2 | 2 | 20 | 724 | 1689 |
| *Slc17a7* | solute carrier<br>family 17<br>(sodium-<br>dependent<br>inorganic<br>phosphate<br>cotransporter),<br>member 7 | Mus<br>musculus | NM_182993.2 | 416631-C2 | 2 | 20 | 464 | 1415 |
| *Slc17a8* | solute carrier<br>family 17<br>(sodium-<br>dependent<br>inorganic<br>phosphate<br>cotransporter),<br>member 8 | Mus<br>musculus | NM_182959.3 | 431261-C2 | 2 | 20 | 781 | 1695 |

### Tissue and brain-region enrichment of Csmd1 protein (Figure 2A, B; Figure 2 – Figure Supplement 1A, B)

Three p60 C57bl6N mice (Charles River) were deeply anesthetized with isoflurane, cervically dislocated, and decapitated. Brain, heart, liver, lung, spleen, muscle, skin, and testes were harvested and frozen on dry ice. Frozen tissue was homogenized in NP-40 lysis buffer (150 mM NaCl, 50mM Tris pH 7.5, 1% NP-40 [Sigma #18896]) with phosphatase and protease inhibitors for 2 × 10 min at 30hz in a Qiagen Tissue Lyser. Lysates were spun 10 minutes at 15,000 x g to pellet insoluble material, supernatant was removed to a new tube and protein concentration assayed by Bradford (Bio-Rad). From each tissue sample, 20ug of protein was incubated with 4x laemmli buffer (Bio-Rad) and separated for 50 minutes at 200V on 4-15% TGX stain-free protein gels (Bio-Rad) on ice. To minimize aggregation of large proteins, samples were not boiled. To enable comparison amongst tissues of all 3 animals samples were run on the same, 26-well gel. Lysate from a *Csmd1* KO mouse was run as a negative control. After separation, gels were activated for 5 minutes with UV excitation to crosslink the gel’s UV-visible protein dye with separated proteins. Subsequently, gels were transferred for 2 hours on ice at 100V onto PVDF membranes (Millipore), blocked for 1 hour at room temperature in 5% milk in TBST (Tris Buffered Saline with 0.02% Tween-20) followed by overnight incubation in 1:1000 anti Csmd1 antibody (Abcam ab166908) in 5% milk TBST. This antibody was generated against a peptide in the transmembrane region of Csmd1. The blot was washed 3 x in TBST for 15 min each and then total protein was imaged with UV excitation in a Bio-Rad GelDoc. Blots were then incubated with anti-rabbit-HRP for 2 hrs. at RT in 5% milk TBST, then washed 3x again in TBST. Bands were visualized with Femto-ECL (Pierce) chemiluminescence, avoiding saturation in any pixels. For quantification, Csmd1 band intensity was normalized to total protein in the same lane using Image J. Statistical comparisons were made with ANOVA with post-hoc Tukey test, correcting for multiple comparisons, in GraphPad Prism. For brain-region enrichment experiments (Figure 5), a similar protocol to above was followed with a separate cohort of 3 WT mice except that brains were solubilized in SDS lysis buffer, protein estimated with BCA assay (Thermo Fisher) and 10 μg total protein separated on 4-15% TGX stain-free protein gels (Bio-Rad). Molecular weight was estimated using HiMark pre-stained protein standard (ThermoFisher) or Precision Plus pre-stained protein standard (Biorad).

### Characterization of *Csmd1* mRNA expression in the retinogeniculate system by Droplet digital RT-PCR (ddRT-PCR) (Figure 2E)

To isolate the LGN, Alexa-488 conjugated CTB was injected under anesthesia into the eyes of p5 mice followed by microdissection under a fluorescent dissecting scope. Retinas and RGCs were retrieved from a pre-prepared tissue bank at −80°C (prepared as in (Barres et al., 1988)). N = 3 for p5 retina. N = 10 p5 LGN, N = 5 RGC preps. RNA was extracted from frozen tissue with Qiagen RNeasy Lipid Micro with on-column DNase digestion according to the manufacturer’s protocol. Ten ng of RNA was used as input for ddRT-PCR (Bio-Rad) according to the manufacturer’s protocol with the below assays. Samples were assayed in technical duplicates. Droplets were counted and ratios of *Csmd1*/*Eif4h* were quantified using Quantasoft software (Bio-Rad).

For *Csmd1* assays, a primer-probe set was designed across exon 1 and 2 of mouse *Csmd1*, encompassing the small region deleted in the KO using Primer 3 software (Koressaar and Remm, 2007; Untergasser et al., 2012) and ordered from Integrated DNA Technology.

***Csmd1***

LEFT PRIMER GGCTCCTCACTGCAGCTAAG
RIGHT PRIMER AAAGCCAGGACTTTCAATGGTG
Probe (5’ FAM, 3’ ZEN quencher) TGTACCAGGCCACCACAGTTCTGACC
Product size: 80bp

The control assay to *Eif4h* consisted of the following:

***Eif4h* (control assay)**
*Eif4h*_mouse_exp_ctrl_F: GTGCAGCTTGCTTGGTAGc
*Eif4h*_mouse_exp_ctrl_R: GTAAATTGCCGAGACCTTGC
*Eif4h*_ctrl_VIC (5’ VIC, 3’ Zen): agcctACCCCTTGGCTCGGG

Thermocycling conditions were as follows:

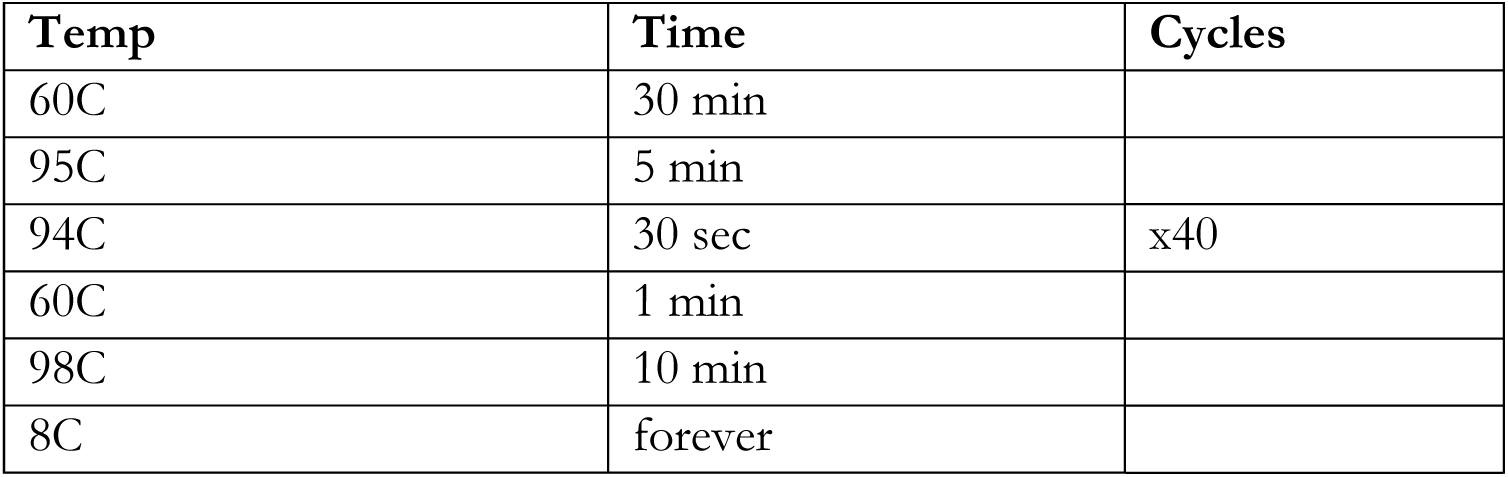

### Csmd1 Immunohistochemistry (Figure 2C, F)

P10 C57Bl6N WT and *Csmd1* KO mice were deeply anesthetized with Avertin (240 mg/kg, i.p.) and transcardially perfused with 20 ml of PBS followed by 20 ml of 4% PFA in PBS. Brains were extracted and drop-fixed for 2hrs on ice in 4% PFA in PBS, washed in PBS, and cryoprotected in 30% sucrose – PBS solution approximately 24-48 hours (until the tissue sank to the bottom of the tube). Brains were embedded in 2:1, 30% sucrose-PBS : OCT (VWR), and stored at −80°C. Cryosections (14 µm) were cut on a Leica cryostat and affixed to Leica Surgipath X-tra slides and processed for IHC as follows. Slides were baked for 20 min at 37°C followed by 3x rinses in PBS. Slides were blocked for 1hr in blocking solution (antibody buffer with 5% BSA and 0.8% Triton X-100) followed by incubation overnight at 4°C with 1:1000 rabbit anti-Csmd1 antibody (Lund, produced by Agrisera, Sweden) in 4:1 antibody buffer (150 mM NaCl, 50 mM Tris Base pH 7.4, 1% BSA, 100mM L-Lysine, 0.04% azide): blocking solution. Anti-Csmd1 antibody (Lund) is a custom polyclonal antibody against the second N-terminal sushi domain of CSMD1 as detailed in Escudero-Esparza et al., 2013. Slides were then washed 3 × 15 min in PBS followed by incubation with 1:200 donkey anti-rabbit IgG Alexa-594 in 4:1 antibody buffer: blocking buffer. Slides were washed 3 × 15 min in PBS, mounted with Vectashield with DAPI (Vector Labs), and sealed with nail polish. Using a Zeiss AxioImager Z1, mosaics were taken with a 10x objective and stitched together or, for high-resolution images, taken as single z-planes at 63x on a LSM700 confocal microscope. Figure 2F was generated through export of the dLGN as a region of interest from the coronal slice mosaic shown in Figure 2C.

### Specificity of anti-CSMD1 antibody by ICC in HEK cells (Figure 2 - Figure Supplement 1C)

HEK 293 cells (ATCC ref CRL-1573) were transfected using Lipofectamine 2000 (Invitrogen 11668-019) and 2 μg of pcDNA3.1 CMV V5 *Csmd1* (*Mus Musculus*) or without DNA (no template control). 24h later cells were fixed using 4% paraformaldehyde and blocked with 10% goat serum, then incubated overnight at 4 degrees Celsius with Anti Csmd1 (1:1000, Lund). Cells were washed and then stained with Alexa 488-conjugated secondary antibody (1:500) at 1/500 for 30 min at room temperature. Cells were washed and visualized on a Zeiss axio imager Z.1. *Csmd1* construct was custom synthesized (Genscript) with N-terminal V5 tag inserted by mutagenesis.

### Quantification of Csmd1 colocalization with synaptic markers (Figure 2 G, H; Figure 2 – Figure Supplement 1 D-F; Figure 4 H, I)

Tissue was prepared and IHC performed as above with the following modifications. The primary antibodies used were (Figure 2) 1:1000 rabbit anti-CSMD1 (Lund) and 1:1000 guinea pig anti-vGluT2 (synaptic systems) or (Figure 4) 1:1000 rabbit anti-CSMD1 (Lund), 1:500 chicken anti vGat (synaptic systems), and a mix of 1:10,000 guinea pig anti vGluT2 (synaptic systems) and of 1:10,000 guinea pig anti vGluT1 (synaptic systems). For analysis, (Figure 2) 8 fields of view per dLGN (4 in the core region and 4 in the shell region) or (Figure 4) 5 FOVs per animal in somatosensory cortex layers II/III were acquired as single z-planes at 63x on an LSM700 confocal microscope. Animals were (Figure 2) 6 or (Figure 4) three p10 WT C57bl6N-Tac mice. Images were processed in custom CellProfiler pipeline to count colocalized puncta like above. As a control for amount of colocalization that would be expected by chance, one channel was rotated by 90 degrees prior to colocalization analysis. Statistical tests compared Csmd1 colocalization with each synaptic marker to this chance value.

### Complement Component C3b deposition on retinogeniculate synapses (Figure 2 I, J)

P10 C57Bl6N WT and *Csmd1* KO littermates were deeply anesthetized with Avertin and transcardially perfused with 20ml of PBS followed by 20ml 4% PFA in PBS. Brains were extracted and drop-fixed 2hrs on ice in 4% PFA in PBS, washed in PBS, and cryoprotected in 30% sucrose – PBS solution approximately 24-48 hours (until the tissue sank to the bottom of the tube). Brains were embedded in 2:1, 30% sucrose-PBS: OCT (VWR), and stored at −80°C. Fresh 14 μm cryosections were cut on a Leica cryostat and affixed to Leica Surgipath X-tra slides and processed the same-day for IHC; using only newly cut, same-day processed slices gave the most reliable C3 staining and littermate pairs were always processed on the same day. Staining and quantification was performed as previously described in (Hong et al., 2016) with modifications. Slides were baked for 20min at 37C followed by 3x rinses in PBS. Slides were blocked 1hr in blocking solution (Antibody buffer as above, with 5% BSA and 0.4% triton X-100) followed by incubation overnight at 4°C with 1:500 rabbit anti-C3 antibody (Dako) and 1:10,000 guinea pig anti-vGluT2 (Synaptic Systems) in 4:1 antibody buffer: blocking solution. Slides were washed 3 × 15 min in PBS followed by incubation with 1:200 donkey anti-rabbit IgG 594, and 1:200 donkey anti-guinea pig IgG 488 in 4:1 antibody buffer: blocking buffer. Slides were washed 3 × 15 min in PBS, mounted with Vectashield with DAPI (Vector Labs) and sealed with nail polish. In imaging, 3-5 single z-planes were captured per dLGN, avoiding the shell where vGluT2 inputs increase in density and brightness. Z-planes were captured at a depth at which C3 and vGluT2 staining was uniform. Images were acquired with identical parameters between each WT and KO pair on a Zeiss LSM 700 confocal microscope. Puncta number and colocalization were quantified using a custom pipeline in CellProfiler based on thresholding the images into binary and identifying overlapping areas. Because there was batch to batch variation in staining intensity between each pair of littermates causing some weightings of the automatic thresholding to be too stringent for some pairs and too lenient for others, each littermate pair was processed as an individual unit for which the stringency of weighting of the automated thresholding algorithm was empirically determined for a test image from one animal to approximate puncta that would be retained by an experienced individual thresholding manually and then applied to all images from both animals of the pair. Results were normalized to the WT in each litter; where there was more than one WT in the pair, the animals were all normalized to the average of the WTs.

### Quantification of retinogeniculate synapses (Figure 3 C, D; Figure 3 – Supplement 2)

Tissue was prepared and IHC performed as above with the following modifications. The primary antibodies used were 1:500 anti-Homer 2 and 1:1000 guinea pig anti-vGluT2. Images captured on an LSM880 confocal microscope with a 63x objective lens consisted of 3-Z-planes 1 µm apart, 8 fields of view per dLGN (4 in the core region and 4 in the shell region). Each plane was analyzed in a custom CellProfiler pipeline to count colocalized puncta; average puncta counts were normalized to WT littermates as above. The puncta counts per animal are the average counts of the z-planes from all FOVs and LGNs for that animal.

### Eye Segregation in the dLGN (Figure 3 A, B; Figure 3 – Figure Supplement 1C, D)

Eye segregation was performed and analyzed with the multi-threshold method as previously described (Lehrman et al., 2018; Stevens et al., 2007). Briefly, under anesthesia, the eyelids of p9 pups were surgically opened, a hole created at the interface of the sclera with a 30-gauge needle, and then 1 μl of either Alexa-594-conjugated or Alexa-488-conjugated cholera toxin beta subunit (CTB) was injected with a blunt-end Hamilton syringe, such that each eye received a different color. Brains were harvested at p10 and fixed overnight in 4% PFA in PBS, then transferred to 30% sucrose PBS for cryoprotection. Brains were sectioned on a frozen microtome at 40 μm thickness and collected into PBS in 24 well plates. Sections containing the dLGN were spread onto VWR Super Frost Plus slides, allowed to airdry, mounted in Vectashield with DAPI, and sealed with nail polish.

dLGNs were imaged on an E800 microscope with a SPOT camera at 10x magnification. To best capture the structure of the fibers in the dLGN while allowing for small differences in fluorescent intensity, each image was captured with an autoexposure. Five sections of the medial-most portions of the LGN were selected per dLGN as input for analysis. In these sections, the ipsilateral patch is located medially in a bullet-shaped dLGN. Images were thresholded manually to best capture LGN fibers and then input into a custom CellProfiler pipeline. A user-guided ROI was drawn around each dLGN to separate it from the surrounding structures like the optic tract, IGL, and vLGN and then the area occupied by the ipsi and contra was quantified, and the size of the contra territory, the ipsi territory, and their overlap was calculated. The overlap percentage was calculated as the area of ipsi/contra overlap divided by the area occupied by the dLGN as defined by the ROI. By convention, each LGN was counted as an “N” in subsequent analysis yielding 2 LGNs per animal. The analysis followed the multi-threshold method designed to examine the signal to noise landscape by varying the threshold stringency of the ipsilateral territory in increments of 5; Unpaired t-tests were used at each threshold as previously described (Stevens et al., 2007; Lehrman et al. 2018). Because the curve of “overlap as a function of threshold-adjustment” risks artifactually flattening at higher stringencies in low-overlap phenotypes (e.g. if increasing the threshold stringency results in local “gaps” between parts of binary areas, further increasing threshold-stringency could not further decrease overlap in that local area), adjustments to make the baseline threshold more lenient (negative % adjustments, i.e. −10, −5%) were included as well as the set of adjustments to make the baseline threshold more stringent (positive % adjustment, i.e. +5, 10, 15, 20, 25, 30%) conventionally used to study phenotypes of high-overlap (e.g. in Stevens et al., 2007).

Quality control occurred at 3 steps. 1) The superior colliculi were examined for full, bright, uniform fill of each color on a fluorescent dissecting microscope; those that lacked a color or had a hypo-intense area were discarded. 2) During slicing and spreading, any animal in which a slice was lost in the LGN was discarded. 3) After imaging, dLGNs were inspected for poor fills, defined as hypo-intense fluorescent areas (or areas with no fluorescence) present in multiple slices that did not geographically correspond to fluorescence in the other color; these hypo-intense areas tended to migrate in the opposite direction of the ipsi patch as one moved through slices in the LGN. If an animal had a spotty fill evident in either color, the whole animal failed quality control.

### Retinal Ganglion Cell Counts (Figure 3 – Figure Supplement 1A, B)

Retinal whole mounts and RGC counts were performed as in (Lehrman et al., 2018) with modifications. Briefly, retinas of p10 WT and KO littermates were isolated and stained free floating with RBPMS. Whole mount retinal preparations were performed by dissecting out the retina of P10, 4% PFA perfused mice (1 retina per animal, n=6 knockout animals, n=14 wildtype animals). Retinas were blocked in staining buffer (10% Normal Goat Serum and 2% Triton X-100 in PBS) for 1 hour before incubation with guinea pig Rbpms (1:500, HM0401, HuaBio) at 4°C for three days. After three 10 minute washes in PBS, retinas were incubated in a secondary antibody (1:200, #1924784, Life Technologies) at room temperature for one hour. Retinas were washed for 10 minutes three times and mounted using glycerol. One retina per animal was imaged in Z-stacks on an LSM 880 confocal microscope to enable accurate cell counting despite the tissue not lying flat. A total of 20 images per animal (12 peripheral retinal images and 8 central retina images) were captured and, blind to genotype, manually scored by number of RBPMS positive cells in ImageJ software (NIH) via the cell counter plugin.

### Developmental profiling in Synaptosome fractions and brain (Figure 4G, Figure 4 – Figure Supplement 1)

Synaptosomes were prepared as previously described (Ehlers et al., 1998). Briefly, 6 C57bl6N (Charles River) animals at each of 4 ages (p5, p10, p30, p60) were deeply anesthetized with isoflurane followed by cervical dislocation and decapitation; there were 3 male and 3 female animals at each age except p5, which had 4 male and 2 female animals. Brains were extracted, and cerebellum and brainstem removed. Forebrains were finely chopped and then homogenized in 0.32 M sucrose solution (0.32 M sucrose, 10 mM Hepes pH 7.4, with cOmplete protease (Roche) and phosphatase inhibitors) using a motorized glass pestle Dounce homogenizer (40 strokes at speed 80). An aliquot of this suspension was kept as total brain homogenate. Samples were spun at 1,200 x g for 10 minutes at 4°C to pellet the nuclear fraction (P1). The supernatant (S1) was collected to a new tube and centrifuged 15,000 x g. The supernatant (S2) was discarded and the pellet (P2) containing the crude synaptosomes was resuspended and processed for western blotting in NP-40 lysis buffer as above with the following modifications: equal amounts of protein per sample (unboiled) were separated on 3-8% Tris-Acetate XT gel (Bio-Rad) in XT Tricine running buffer (Bio-Rad) and samples were normalized to β-Actin (Santa Cruz sc-47778). Molecular weight was estimated using Invitrogen MagicMark XP.

### Engulfment of synaptosomes by cultured microglia (Figure 5 A-D, Figure 5 – Figure Supplement 1 A-E)

Crude synaptosomes were prepared as in (Ehlers et al., 1998) with modifications. Animals were processed in batches of 6 mice, with littermates always being processed together. Forebrains from 6-week old sex-matched WT-*Csmd1* KO littermates were finely chopped and then homogenized in 0.32 M sucrose solution (0.32 M sucrose, 10mM Hepes pH 7.4, with complete protease and phosphatase inhibitors). Samples were spun at 1,200 x g for 10 minutes at 4°C to pellet the nuclear fraction (P1). The supernatant (S1) was collected to a new tube and centrifuged at 15,000 x g for 15 minutes to pellet down synaptosome fraction. The supernatant (S2) was discarded and the pellet (P2) containing the crude synaptosomes was resuspended in 0.1M sodium bicarbonate (pH 8.4) and mixed with Alexa 647 dye (Alexa Fluor 647 NHS Ester) at a concentration of 10ul dye stock/1mg protein and rocked at room temperature for 1 hour. Labeled synaptosomes were centrifuged at 15000g for 3 minutes and washed with PBS 4 times. After final wash, labeled synaptosomes were resuspended in 0.32 M sucrose with 5% DMSO and stored at −80°C.

Primary mouse microglia were prepared from P15 C57Bl6J mice. Microglia isolation was performed as described in (Pino and Cardona, 2011) with modifications. Briefly, mice were transcardially perfused with cold HBSS and brains with cerebellum removed were manually homogenized in RPMI. Homogenate was applied to a Percoll gradient and centrifuged at 500g for 30 minutes at 4°C. Cells were collected from the 30-70% interface, pelleted, washed with HBSS, and resuspended in culture medium (DMEM/F-12 + GlutaMAX, 10% fetal bovine serum, 1X Penicillin-Streptomycin, 20ng/ml mCSF).) Microglia were then cultured in 96-well format (10Kcell/well) for 4-5 days. On the day of the assay, labeled synaptosomes were resuspended in DMEM media to achieve an optical density of 0.8 at absorbance 600 nm in order to normalize for synaptosome concentration. Synaptosome solution was added to cells at a final synaptosome solution to media concentration of 5:200. At 15, 30, or 45min post synaptosome addition (according to the desired time of data collection), cells were washed 3 times with PBS and then fixed with 4% paraformaldehyde and stained with anti-Iba antibody (Fuji Film Wako 019-19741, 1:500). Cells were imaged and analyzed using a Thermo Fisher Arrayscan XTi. A 20X objective was used to image 64 fields of view per well. Cell masks were generated to delineate individual cells and total Alexa 647 fluorescence intensity per cell was measured and used as a reflection of amount of engulfed material.

---

**Figure 1 – Figure Supplement 1.**
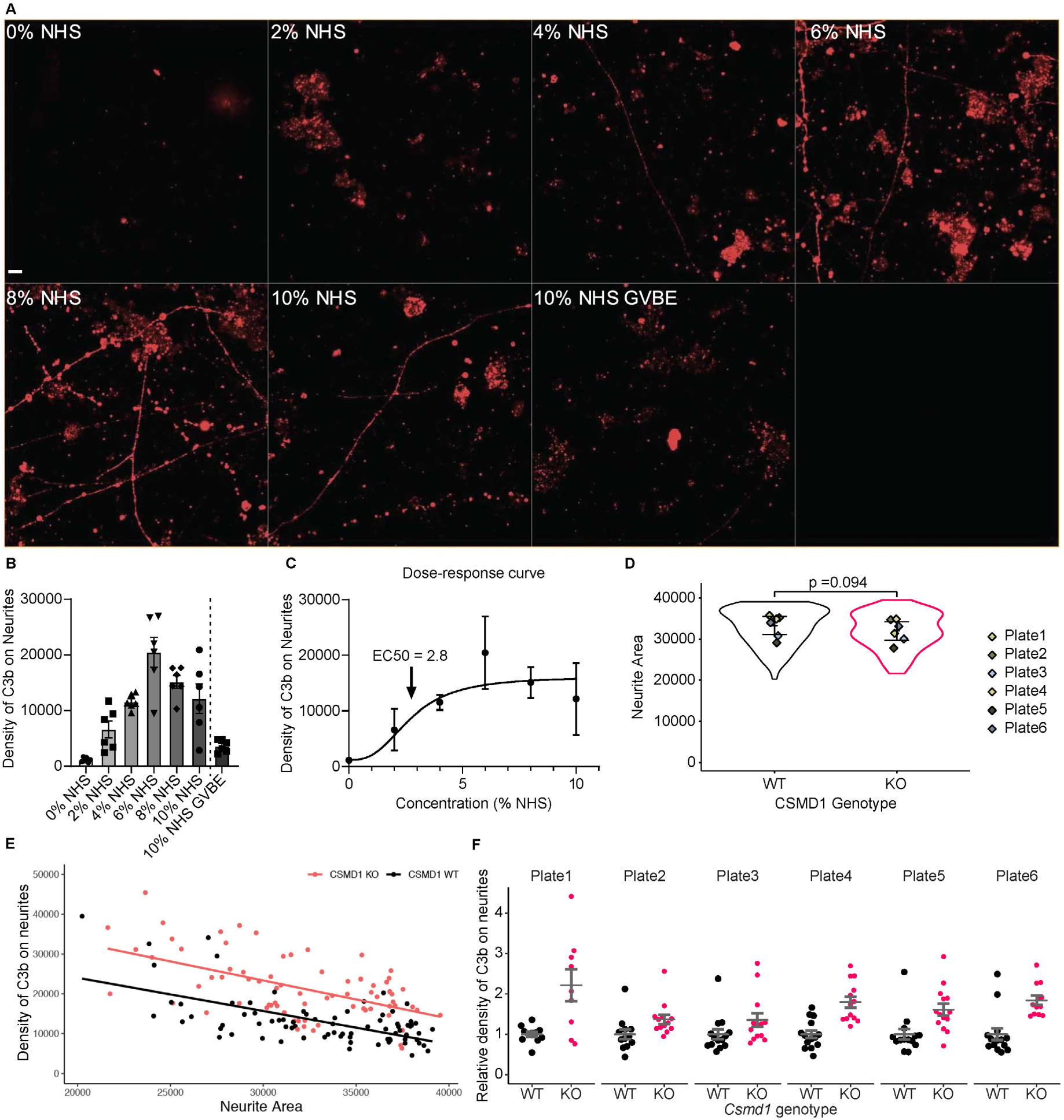
The complement deposition assay was performed within the dynamic range of NHS-complement deposition dose-response curve, and genotype effect was robust to cell density and across plates. A) Representative image from experiment to establish a NHS dose-response curve in the complement deposition assay; Scale = 10um. Briefly, at day 14, differentiated neurons were primed to activate complement through the generation of surface immune complexes, then challenged with different doses of normal human serum as a source of complement; the EDTA-containing buffer GVBE was used as a negative control, as the EDTA inactivates the convertases of the complement cascade. Extent of C3b deposition was quantified by live staining, followed by fixation and imaging on a Perkin Elmer Opera Phenix high content confocal imager with a 63x objective. C3b signal alone in wells of WT neurons is shown (red/orange); notice that neuritic C3b-labeling emerged and increased with serum concentration but was not seen in the GVBE control. Conditions quantified in B) with n= 6 wells each condition across a single plate, where each well represents the average of all FOVs in that well. C) 3% NHS was chosen for the assay in Fig 1 as it lay in the middle of the dose-response curve based on the EC50 of 2.8% calculated by fitting a sigmoid to the data to generate a dose-response curve; a 4-parameter variable slope sigmoid model, Y=Bottom + (X^Hillslope)*(Top-Bottom)/(X^HillSlope + EC50^HillSlope), was fit using the likelihood ratio asymmetric method with robust fit to minimize the effect of outliers (graphpad Prism); error bars are SD. D) Data summarized in Fig 1E re-plotted to show that local neurite area does not differ significantly across genotypes when accounting for plate-level variation. E) Non-normalized data from the Fig 1E assay reveals that complement deposition is correlated with local neurite area for both CSMD1 WT and KO neurons and that CSMD1 KO neurons have more C3 deposition when controlling for this effect of neurite area variation. Analysis of covariance (ANCOVA) was performed to analyze the effect of CSMD1 KO or WT genotype while controlling for variation in GFP-positive neurite area. The slopes of the regression lines for WT and KO neurons are not significantly different (F(1,150)=0.346, P= 0.55). When controlling for neurite area variation, KO CSMD1 neurons have significantly more C3 deposition onto neurites than WT (F(1,150)=84.366, P < 3.13×10^-16). F) Data summarized in Fig 1E re-plotted as individual plates.

**Figure 2 – Figure Supplement 1.**
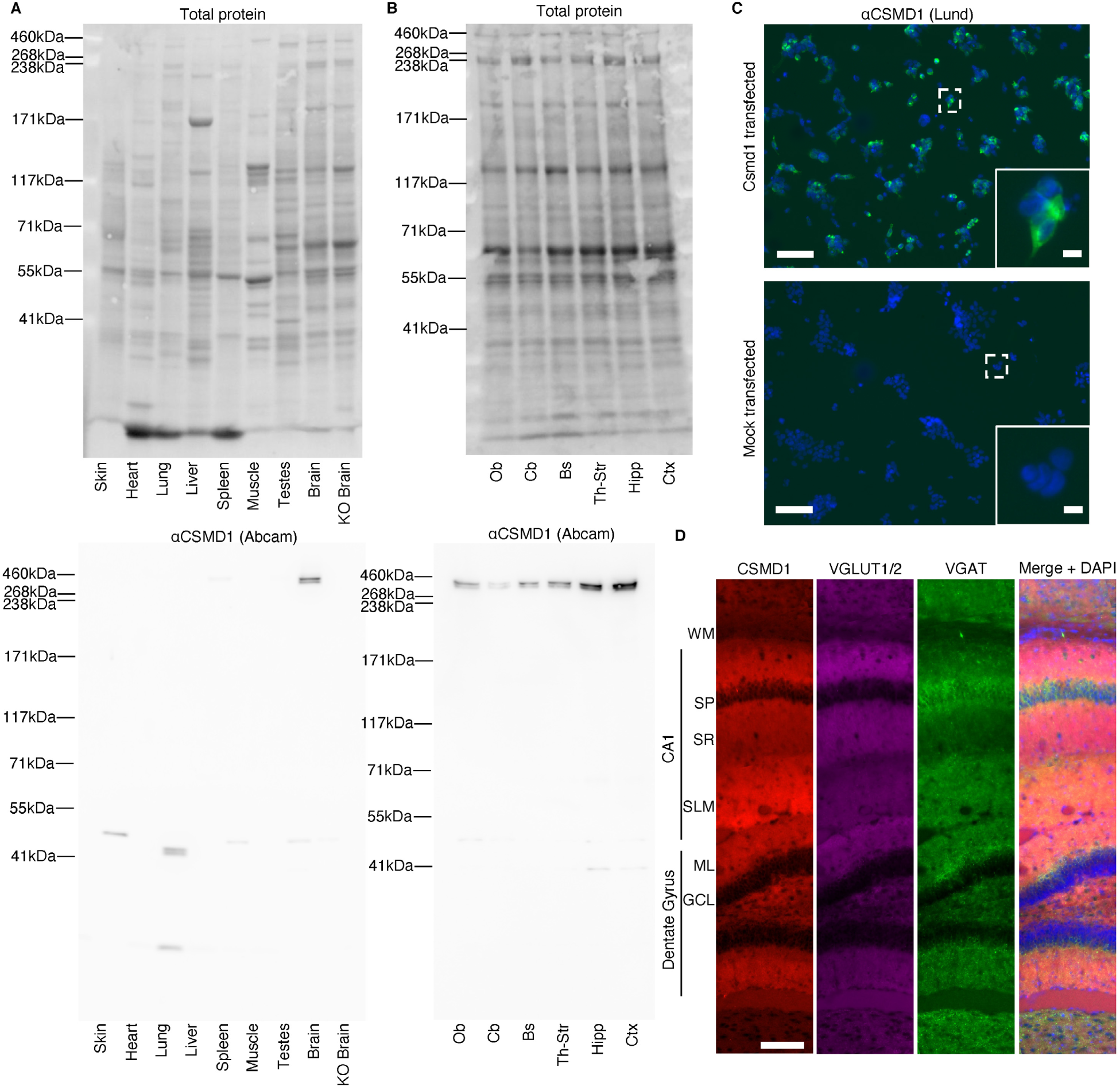
A), B) Top: Total protein blot (Bio-Rad stain-free technology) used to normalize the western blots of Csmd1 in tissue types (Figure 2A) and brain regions (Figure 2B) respectively; Bottom: Full-length lanes of the chemiluminescent anti-CSMD1 signal (∼400 kDa doublet band); approximate molecular weights are shown, which were estimated with HiMark Pre-stained Protein standard (thermofisher). Specificity of anti-Csmd1 antibody (Lund Custom made by Agrisera, Sweden). HEK cells were transfected with either a mouse Csmd1 expression construct (upper panel, *Csmd1 pcDNA3.1*) or empty vector (lower panel, Mock Transfected). Immunocytochemistry with the polyclonal rabbit anti-Csmd1 antibody from Lund (1:1000) yielded signal (green) only in the transfected cells, further suggesting the nuclear signal seen when staining mouse tissue is non-specific. Scale bar overview image = 100 μm, inset = 10 μm. D) Higher resolution image showing CSMD1 immunoreactivity present in synaptic layers of hippocampus and qualitatively low in the overlying white matter tract; scale bar = 100 µm; WM white matter, sp stratum pyramidale, sr stratum radiatum, slm stratum lacunosum moleculare, ml molecular layer, gcl granule cell layer; images are tiled field of views on a conventional fluorescence microscope from a region of interest of the brain slice in Fig. 2 C.

**Figure 2 – Figure Supplement 2.**
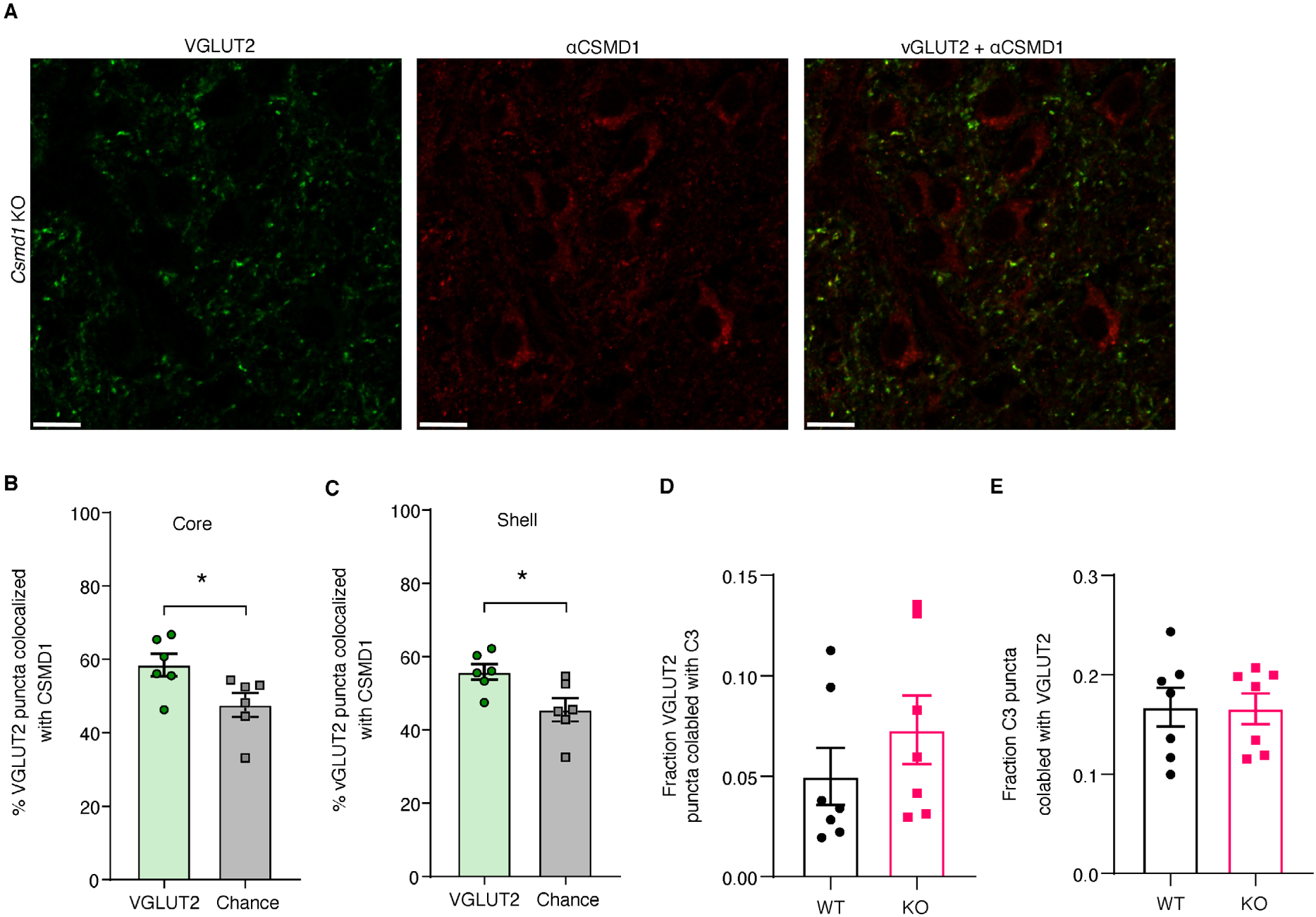
A) Staining of *Csmd1* KO LGN with anti-Csmd1 antibody confirms that punctate neuropil staining in WT LGN at 63x confocal resolution is specific for Csmd1. Scale bar = 10 μm. B,C) Csmd1 puncta colocalize with vGluT2 puncta in both the core (B) and shell (C) regions of the dLGN at a rate greater than chance (rotation control); * p < 0.05, unpaired two-tailed t-test. D,E) Fraction of VGLUT2 puncta co-labeled with C3b (D) and fraction of C3b puncta co-labeled with VGLUT2 (E) are shown here to accompany the normalized statistical analysis of C3b deposition on VGLUT2 terminals in Fig 2. Each data point is the average of FOVs in one animal; Error bars SEM; no statistical comparison was performed.

**Figure 3 – Figure Supplement 1.**
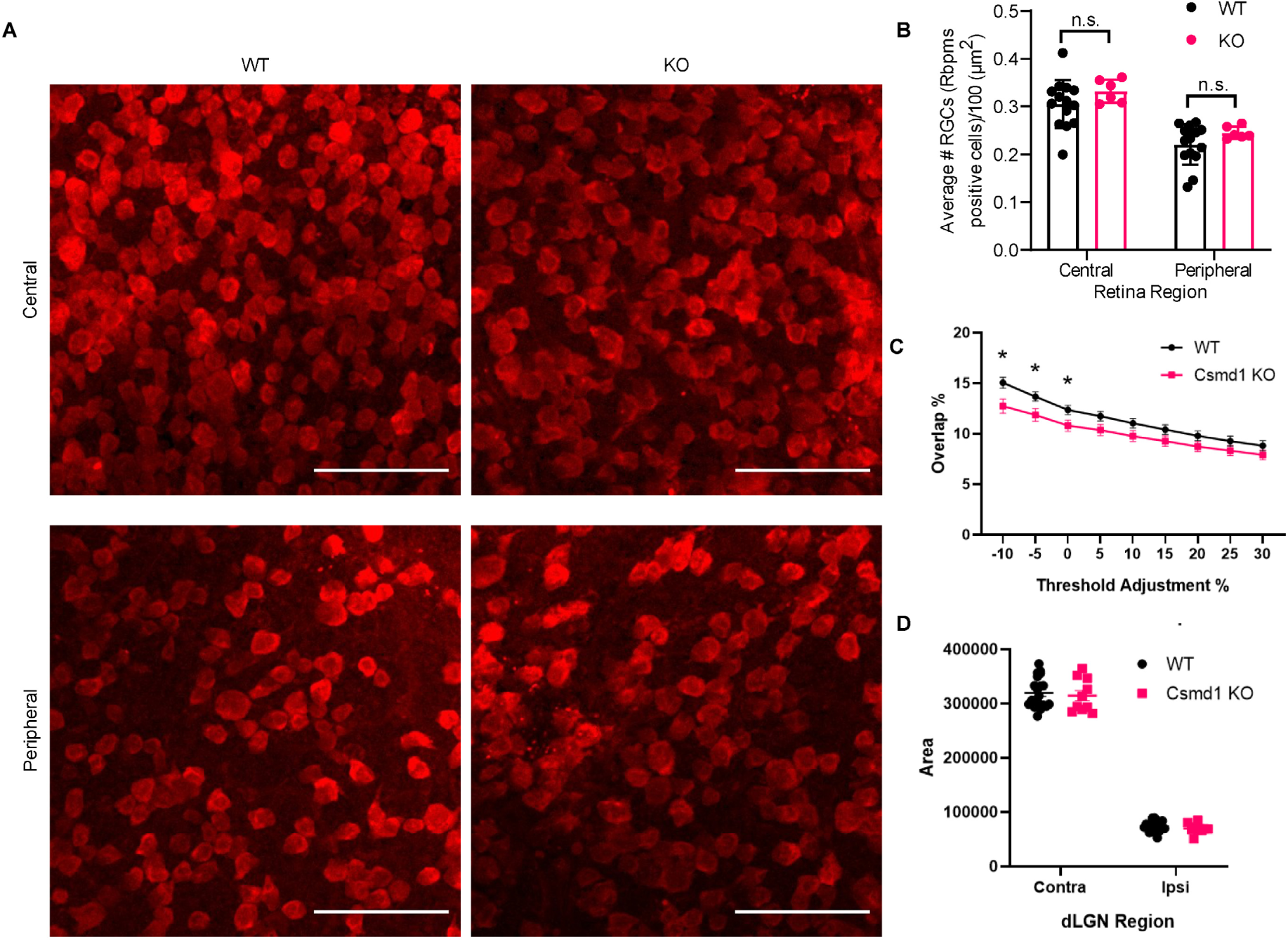
Retinal Geniculate Cell (RGC) numbers and dLGN sizes are not significantly different between WT and Csmd1 KO animals at p10. A) Representative images from central (top) and peripheral (bottom) retina in retinal whole mounts of WT and *Csmd1* KO p10 animals; images are maximum intensity projections of full-thickness z-stacks; scale bar = 75 μm. B) Quantification of number of RGCs in WT and KO central and peripheral retina; by two-way ANOVA mixed-effects model, there was a significant effect of anatomical region (p<0.0001) but not of genotype or anatomical region x genotype (p > 0.05); RGC counts of WT vs KO in each region were non-significant (n.s., p>0.05) by Sidak’s multiple comparisons test; n = 14 WT animals and 6 KO animals. C) Eye-segregation overlap % from figure 3H displayed in a more traditional line plot. * p < 0.05, unpaired 2-tailed t-test; Error bars are SEM. D) Area of the contralateral and ipsilateral dLGN is not significantly different WT compared to KO at p10 in the animals from Fig 3 B, p>0.05 unpaired two-tailed t-test.

**Figure 3 - Figure Supplement 2.**
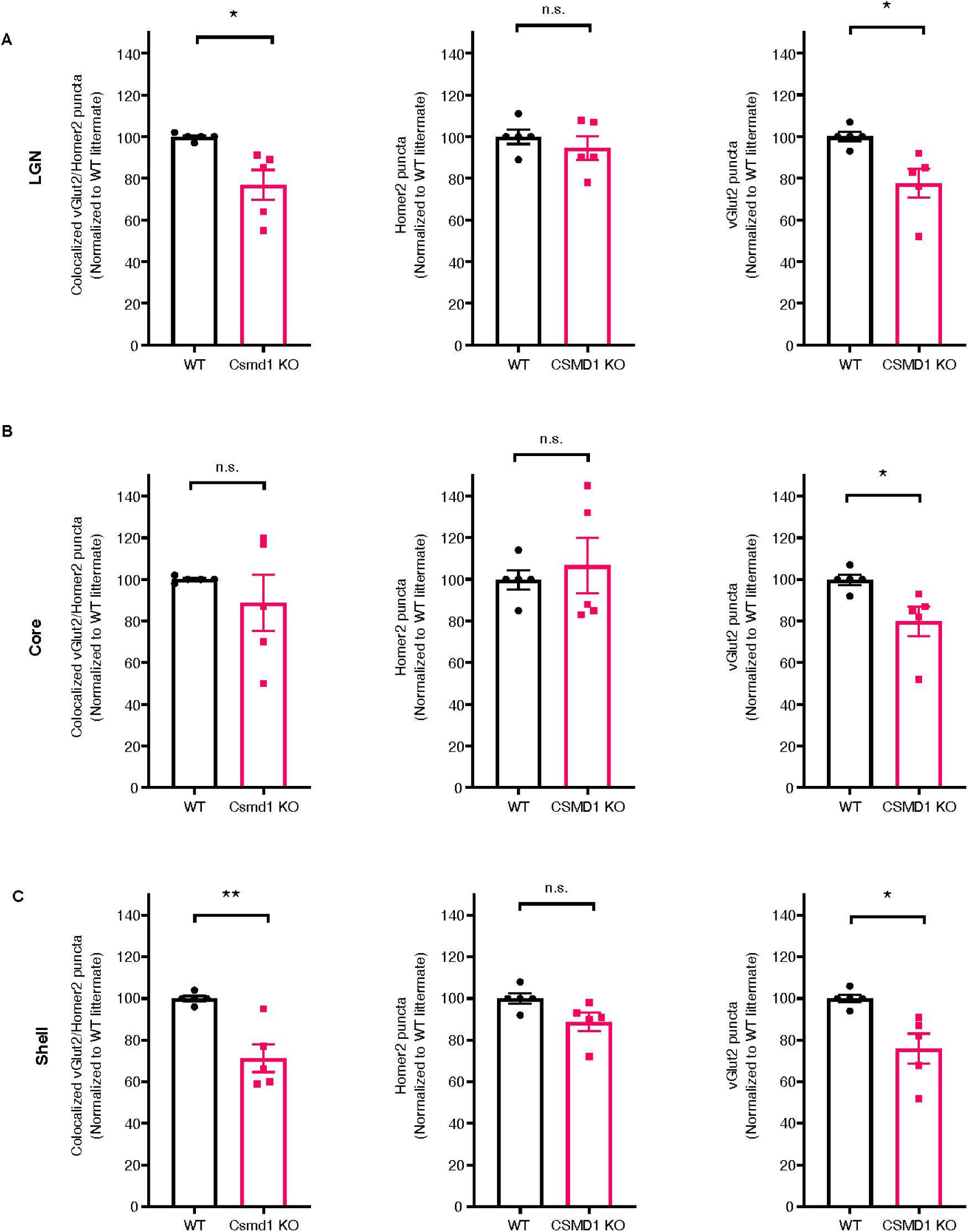
Loss of Csmd1 reduces presynaptic and postsynaptic markers in the dLGN and its sub-regions. At p10, loss of Csmd1 led to reduced retinogeniculate synapses (colocalized vGluT2/Homer2) when averaged across the whole dLGN (A) and the “shell” sub-region (C). Loss of Csmd1 reduced vGluT2 puncta across the dLGN (A), and in the “core” (B) and “shell” (C) sub-regions. Puncta were normalized as a percentage of the value of the WT(s) of the littermate pairs; n=5 littermate pairs, *p<0.05, **p<0.01 unpaired two-tailed T-test.

**Figure 4 – Figure Supplement 1:**
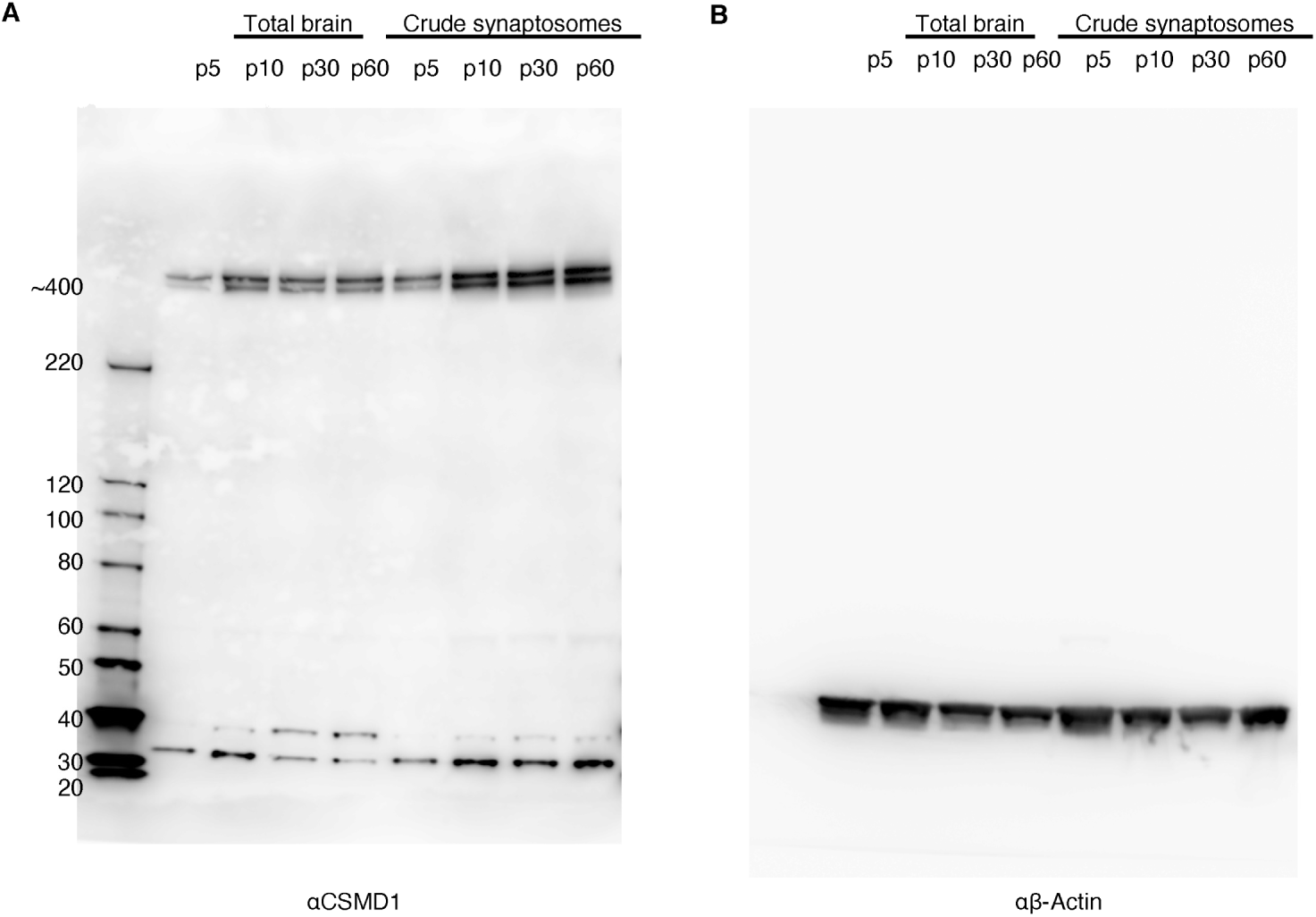
Detail of western blots. Full length lanes of Western Blot of Csmd1 (A) and B-actin (B) in brain lysates and synaptosomes corresponding to Fig 4 panel A. The ∼400 kda doublet represents full-length Csmd1.

**Figure 5 – Figure Supplement 1:**
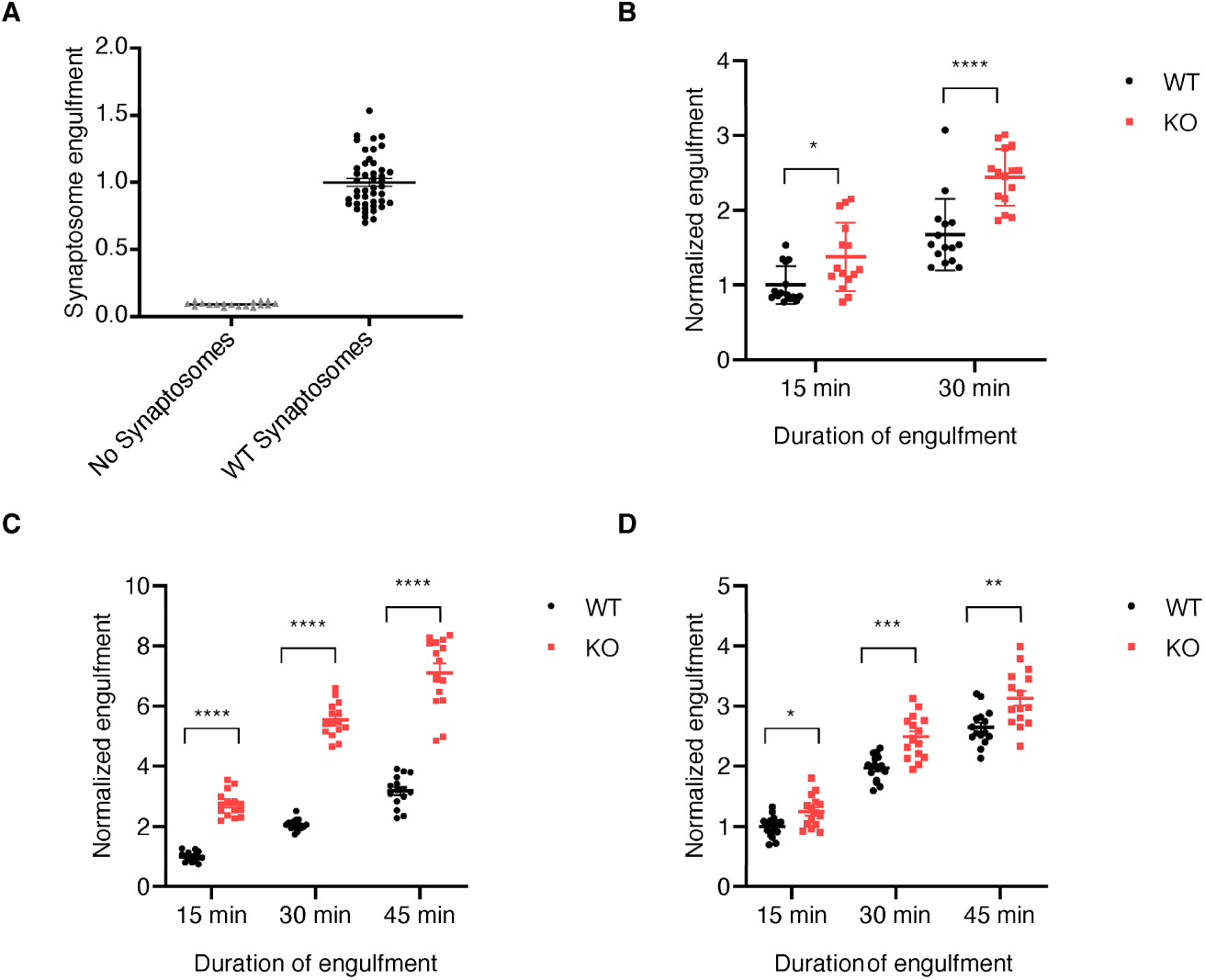
individual experiment data for the *in vitro* engulfment of *Csmd1* KO and WT synaptosomes. A) Signal from wells of microglia fed WT synaptosomes is easily distinguished from the noise in wells with microglia not fed synaptosomes; each data point is a well normalized to the average of the WT wells on that plate; n = 18 wells without synaptosomes and 45 wells with WT synaptosomes, pooled from 3 plates at the 15 min time point. B-D) Data from individual experiments displayed pooled in Fig. 5C,D. Microglia engulfed more synaptosome substrate from *Csmd1* KO animals compared to WT at all timepoints assayed in each experiment; n = 15 wells per genotype. * p<0.05, ** p<0.01, *** p<0.001, **** p<0.0001; 2-way ANOVA with Sidak’s multiple comparisons test.

## Acknowledgements

We thank Dane Hazelbaker and Lindy Barrett for providing CSMD1 KO embryonic stem cells; Daniel Weiner, Lasse Dissing Olesen, and Alec Walker for help in early translations of a complement deposition assay to neurons; Nolan Kamitaki and Giulio Genovese for insights on the genetics of *CSMD1*; Heather De Rivera for technical advice on working with human neurons; Kevin Mastro for discussion and help with mouse necropsy; Molly Heller for technical assistance in assay development; Dorathy Vargas and Joel Cuadrado for animal husbandry; Amy Jarvis and Emma Stickgold for administrative support; Christina Usher for help on figure aesthetics. Mike Carroll, Jessy Presumey, Aswin Sekar, Chinfei Chen, Bernardo Sabatini, and Steve Hyman for useful feedback; Emily Lehrman for technical training; members of the Stevens lab, McCarroll Lab, Carroll Lab, and Eggan Lab for helpful discussion; Douglas Richardson and the Harvard Center for Biological Imaging, the Cellular Imaging Core at Boston Children’s Hospital (C. Chen; NIH U54 HD090255), and the Harvard Neurodiscovery Center Enhanced Neuroimaging Core for infrastructure and support; Sabina Berretta and the Harvard Brain Tissue Resource Center (HBTRC; McLean Hospital) for enabling the localization of expression in human tissue.

